# Natural product-mediated reaction hijacking mechanism validates *Plasmodium* aspartyl-tRNA synthetase as an antimalarial drug target

**DOI:** 10.1101/2025.03.20.644283

**Authors:** Nutpakal Ketprasit, Chia-Wei Tai, Vivek Kumar Sharma, Yogavel Manickam, Yogesh Khandokar, Xi Ye, Con Dogovski, David H. Hilko, Craig J. Morton, Anne-Sophie C. Braun, Michael G. Leeming, Bagale Siddharam, Gerald J. Shami, P.I. Pradeepkumar, Santosh Panjikar, Sally-Ann Poulsen, Michael D.W. Griffin, Amit Sharma, Leann Tilley, Stanley C. Xie

**Author notes:** These authors jointly supervised this work (Michael D.W. Griffin, Amit Sharma, Leann Tilley, Stanley C. Xie). For correspondence (AS); (MDWG); (LT); (SCX). These authors contributed equally to this work (Nutpakal Ketprasit, Chia-Wei Tai, Vivek Kumar Sharma).

## Abstract

Malaria poses an enormous threat to human health. With ever-increasing resistance to currently deployed antimalarials, new targets and starting point compounds with novel mechanisms of action need to be identified. Here, we explore the antimalarial activity of the *Streptomyces sp* natural product, 5ʹ-*O*-sulfamoyl-2-chloroadenosine (dealanylascamycin, DACM) and compare it with the synthetic adenosine monophosphate (AMP) mimic, 5-*O*-sulfamoyladenosine (AMS). These nucleoside sulfamates exhibit potent inhibition of *P. falciparum* growth with an efficacy comparable to that of the current front-line antimalarial dihydroartemisinin. Exposure of *P. falciparum* to DACM leads to inhibition of protein translation, driven by eIF2α phosphorylation. We show that DACM targets multiple amino acyl tRNA synthetase (aaRS) targets, including the cytoplasmic aspartyl tRNA synthetase (AspRS). The mechanism involves hijacking of the reaction product, leading to the formation of a tightly bound inhibitory amino acid-sulfamate conjugate. We show that recombinant *P. falciparum* and *P. vivax* AspRS are susceptible to hijacking by DACM and AMS, generating Asp-DACM and Asp-AMS adducts that stabilize these proteins. By contrast, human AspRS appears less susceptible to hijacking. X-ray crystallography reveals that apo *P. vivax* AspRS exhibits a stabilized flipping loop over the active site that is poised to bind substrates. By contrast, human AspRS exhibits disorder in an extended region around the flexible flipping loop as well as in a loop in motif II. These structural differences may underpin the decreased susceptibility of human AspRS to reaction-hijacking by DACM and AMS. Our work reveals *Plasmodium* AspRS as a promising antimalarial target and highlights structural features that underpin differences in the susceptibility of aaRSs to reaction hijacking inhibition.

## Introduction

In 2023, *Plasmodium falciparum* caused 263 million cases of malaria, resulting in 597,000 deaths, mostly of African children [1]. Resistance of the mosquito vectors to pyrethroid insecticides [2] and widespread resistance of parasites to currently used therapies [3] was compounded by disruptions to prevention strategies during the COVID-19 pandemic [4]. The recent emergence of artemisinin resistance-conferring K13 mutations in Africa [5, 6] is of particular concern. Thus, there is a need to explore new avenues for the development of antimalarial compounds with novel mechanisms of action.

Protein translation relies on aminoacyl-tRNA synthetases (aaRSs) to charge tRNAs with their cognate amino acids [7]. The essentiality of this process means aaRSs represent promising potential targets for antimalarials [8, 9]. For example, the natural product-derived mupirocin, an IleRS inhibitor, is widely used as a topical antibiotic [10], while halofuginone, a ProRS inhibitor, is used to prevent coccidiosis in poultry [11].

Nucleoside sulfamates are chemical compounds that use an unusual reaction-hijacking mechanism to inhibit enzyme function, resulting in new clinical candidates (*e.*g. Pevonedostat, TAK-243 and TAK-981 [12–14]). For many years, the reaction hijacking mechanism was thought to be applicable only to ubiquitin-activating (E1) enzymes. Recently, two different classes of AMP-mimicking nucleoside sulfamates/ sulfonamides have been explored that inhibit aaRSs via a reaction-hijacking mechanism. The pyrazolo-pyrimidine sulfamates, ML901 [15] and ML471 [16], and the aminothieno pyrimidine benzene sulfonamide, OSM-S-106 [17], were shown to target *P. falciparum* TyrRS and AsnRS, respectively, providing potent and specific anti-plasmodial activity. The target enzymes catalyze coupling of the nucleoside sulfamate/ sulfonamide to the amino acid substrate, forming a tight binding uncleavable adduct [15–17].

While ML901, ML471 and OSM-S-106 target specific aaRSs, adenosine 5ʹ-sulfamate (AMS), which is a direct bioisostere of adenosine monophosphate (AMP), was shown to be a broadly reactive hijacking inhibitor of both Class I and Class II *P. falciparum* aaRSs [15]. This opens the possibility of designing bespoke nucleoside sulfamates with tunable specificity.

To explore the range of potential hijackable targets, here we investigated a natural analogue of AMS, known as dealanylascamycin (5ʹ-*O*-sulfamoyl-2-chloroadenosine; DACM) (Fig. 1A). DACM is a natural product nucleoside sulfamate from a soil-dwelling *Streptomyces* bacterium. Earlier studies showed that it exhibits broad-spectrum antibacterial and herbicidal activities [18, 19]. The mechanism of inhibition was not elucidated, but inhibition of bacterial protein translation was reported [20–23]. We hypothesized that DACM is likely a reaction-hijacking inhibitor.

**Figure 1.**
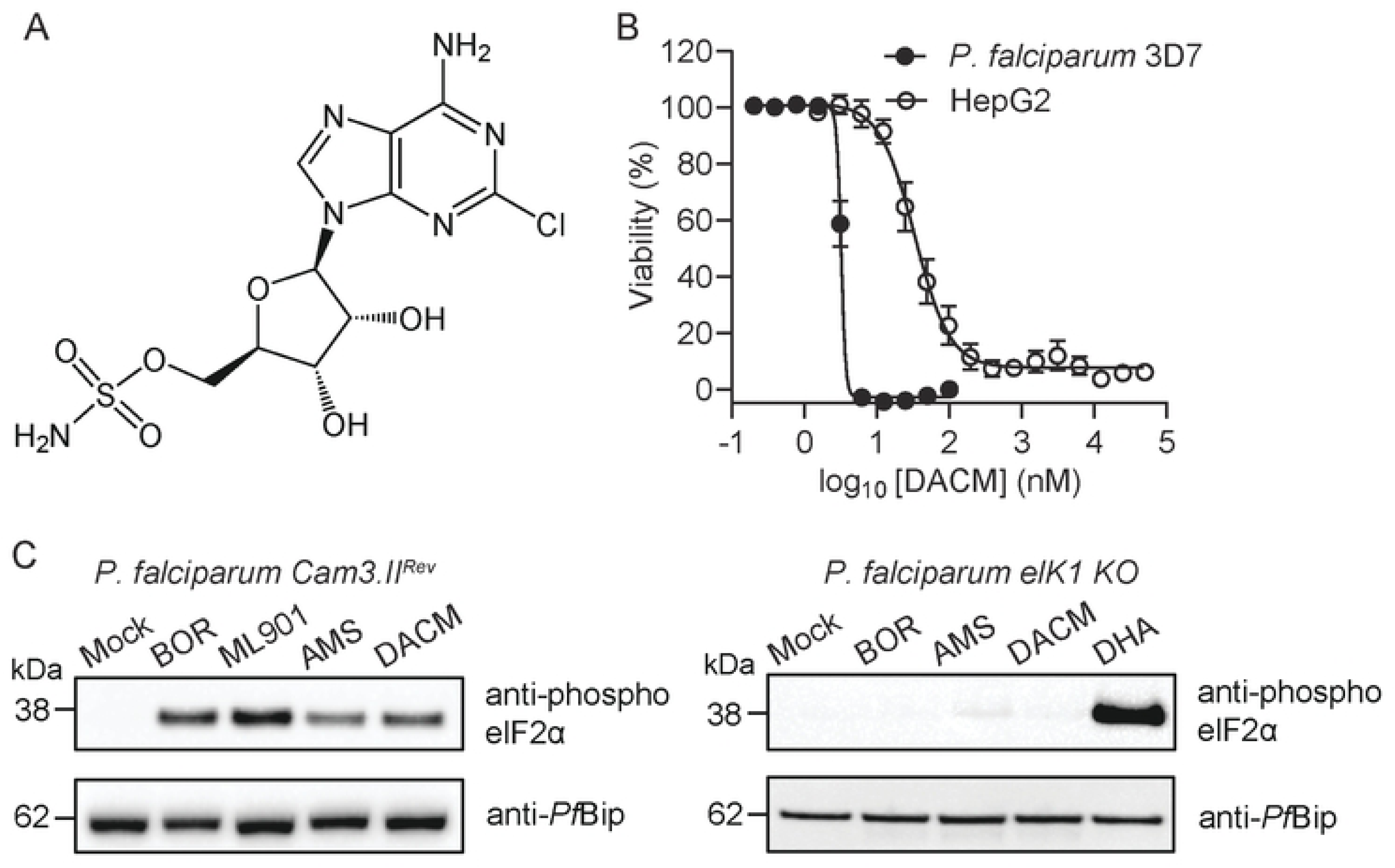
DACM inhibits *P. falciparum* growth and induces the amino acid starvation response. (A) Structure of DACM. (B) Sorbitol-synchronized ring stage parasites were subjected to a 72-h drug sensitivity assay with DACM (white circles). Data represent five independent experiments, each performed in duplicate. Cytotoxicity of DACM (black circles) against the HepG2 mammalian cell line. Data represent four independent experiments each performed in triplicate. Error bars indicate SEM. (C) Trophozoite-stage *P. falciparum* Cam3.II_rev or *P. falciparum* eIK1-knockout cultures were exposed to 200 nM borrelidin (BOR), 1 μM ML901, 1 μM AMS, 1 μM DACM or 1 μM DHA for 3 h. Parasite extracts were analysed by Western blot analysis for phosphorylated eIFα. *Pf*Bip is the loading control. The blot is representative of three independent experiments.

Here we show that DACM inhibits growth of *P. falciparum* cultures by inhibiting protein translation. Targeted identification of DACM adducts confirms multiple aminoacyl tRNA synthetase targets in *P. falciparum*, including cytoplasmic aspartyl tRNA synthetase (AspRS). Via thermal stabilization, targeted mass spectrometry and structural studies of recombinant AspRS from *P. falciparum* and *P. vivax*, we reveal the molecular basis for differential susceptibilities between host and parasite enzymes. This work thus provides a new focus for antimalarial drug development centered on AspRS.

## Results

### Synthesis of DACM, Asp-DACM and Asp-AMS

The synthesis of DACM has been described previously [24, 25]. The synthesis of 5′-*O*-(*N*-L-aspartate)-sulfamoyl-adenosine (**Asp-AMS;** Scheme S1) was performed as detailed in Supplementary Data. 2-chloro-5ʹ-*O*-[*N*-(L-aspartyl)-sulfamoyl] adenosine (Asp-DACM) was prepared by adapting literature methods [26, 27]. Briefly, the bis-protected *N-*(Boc)-*tert*-butyl aspartic acid was converted to the *N-* hydroxysuccinate ester (**1**) with *N*-hydroxysuccinimide and *N*,*N’-*dicyclohexylcarbodiimide (DCC). Next, the aspartate *N-*hydroxysuccinimide ester (**1**) was coupled directly with the 5ʹ-*O*-sulfamoylated isopropylidene protected adenosine (**2**) in the presence of 1,8-diazabicyclo[5.4.0]undec-7-ene (DBU) in *N*,*N*-dimethylformamide (DMF) to yield the triply protected *N-*aminoacylated sulfamoyl adenosine derivative (**3**). Lastly, the complete global deprotection of **3** with trifluoroacetic acid (TFA) in water and tetrahydrofuran (THF) gave the desired 2-chloro-5ʹ-*O*-[*N*-(L-aspartyl)-sulfamoyl] adenosine derivative, **Asp-DACM**, in high purity after precipitation from acetonitrile (MeCN) and triethylamine (Et_3_N) (Scheme 1).

**Scheme 1.** Synthesis of Asp-DACM via coupling of Asp(Boc)-*tert*-butyl ester *N*-hydroxy succinate (**1**) to the 2-chloro-2ʹ,3ʹ-*O*-isopropylidene-5ʹ-*O*-sulfamoyl adenosine (**2)** followed by global deprotection mediated by trifluoroacetic acid (TFA).

### Effect of DACM on P. falciparum growth, stress response and protein translation, and toxicity to mammalian cells

Synchronized ring stage parasites (3D7 strain) were exposed to increasing concentrations of DACM, and parasite viability was assessed in the next cycle by flow cytometry [28]. DACM is a potent inhibitor of the growth of *P. falciparum* cultures (Fig. 1B; IC_50_72h_ = 3.3 ± 0.1 nM) with an efficacy similar to the current front-line drug, dihydroxyartemisinin (DHA) [17]. DACM shows 10-fold reduced toxicity against the mammalian cell line, HepG2, with an IC_50_48h_ value of 47 ± 10 nM (n = 4) (Fig. 1B). Adenosine 5ʹ-sulfamate (AMS, Supp Fig. 1A), a structurally related compound [15], was also highly cytotoxic to *P. falciparum* cultures (Supp Fig. 1B; IC_50_72h_ = 3.8 ± 0.2 (n = 5)), and exhibits a lower level of toxicity against HepG2 (IC_50_48h_ = 53 ± 2 nM (n = 5)) (Supp Fig. 1B).

Accumulation of uncharged tRNAs triggers the amino acid starvation stress response. Using established methods [17], we showed that eukaryotic initiation factor 2α (eIF2α) is phosphorylated upon exposure of *P. falciparum* cultures to DACM and AMS (Fig. 1C, left panel). A similar response was observed for ML901, a known reaction hijacking inhibitor of *Pf*TyrRS, as well as the conventional tRNA synthetase inhibitor, borrelidin (Fig. 1C, left panel). By contrast, in transfectants in which the eukaryotic translation initiation factor 2-alpha kinase 1 (EIK1) has been deleted [29], DACM and other aaRS inhibitors did not cause eIF2α phosphorylation (Fig. 1C, right panel). As previously reported [30], exposure of the EIK1 knockout to dihydroxyartemisinin (DHA) still induces eIF2α phosphorylation (Fig. 1C, right panel).

The aaRSs charge tRNAs with amino acids to drive protein synthesis. We monitored protein translation in *P. falciparum* by following the incorporation of an alkyne analogue of puromycin (OPP), followed by attachment of a clickable fluorophore [31, 32]. Upon treatment of schizont stage parasites with DACM or AMS, protein translation was inhibited with IC_50_3h_ values of 32 ± 1 and 27 ± 6 nM, respectively (Fig. 2A,B). These values correlate well with the IC_50_3h_ values observed for parasite killing of 22.6 ± 0.2 and 18 ± 4 nM, respectively (Fig. 2A,B). These data are consistent with inhibition of protein translation being directly linked to parasite killing. An equivalent 3-h exposure to the folate pathway inhibitor, WR99210, kills parasites without affecting protein translation in the period monitored (Fig. 2C), while a ribosome-directed protein translation inhibitor, cycloheximide, inhibits translation, but this short exposure results in limited killing (Fig. 2D).

**Figure 2.**
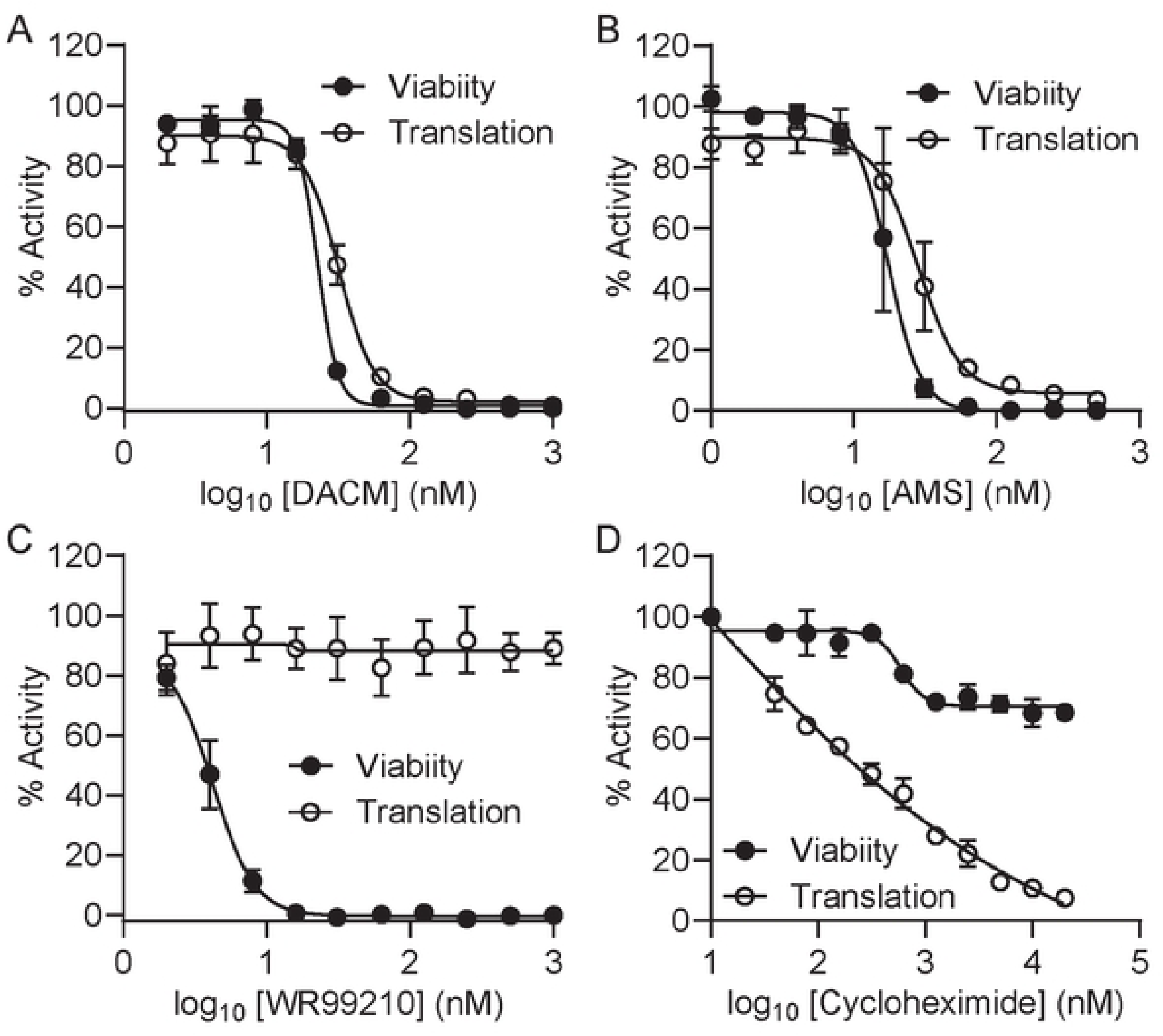
DACM treatment inhibits protein translation in *P. falciparum* cultures. Late-stage *P. falciparum* (Cam3.II_rev) infected RBCs were exposed to the inhibitors for 6 h with the incorporation of OPP during the final 2 h. For parasite viability, compounds were washed out after the 6-h exposure, and viability was assessed in the next cycle. (A) Cycloheximide, (B) AMS, (C) DACM, (D) WR99210.

### DACM targets multiple aaRSs in P. falciparum, including PfAspRS but not PfTyrRS

Reaction hijacking nucleoside sulfamates attack the activated oxy-ester bonds of the enzyme-bound aminoacyl tRNA to form tight-binding adducts (Fig. 3A). We used targeted mass spectrometry to search for potential conjugates in *P. falciparum*-infected red blood cells (RBCs) that had been treated with DACM (10 μM, for 3 h). Following Folch extraction of lysates, the aqueous phase was subjected to LCMS and the anticipated masses for the 20 amino acid conjugates were interrogated. Signals were observed with precursor *m/z* values within 5 ppm of theoretical values calculated for DACM adducts of Asn, Asp, Thr, Ser and Lys, and weaker signals were found for His and Phe (Fig. 3B,C; Supp Fig. 2). The identity of Asp-DACM adduct was confirmed using a synthetic standard, which exhibited essentially the same retention time, precursor ion *m/z* value and MS/MS fragmentation spectrum as the species generated by *P. falciparum* (Fig. 3B-D). No corresponding peaks were detected in control samples not exposed to DACM. The data indicate that at least *Pf*AsnRS, *Pf*AspRS, *Pf*ThrRS, *Pf*SerRS, *Pf*LysRS, *Pf*HisRS and *Pf*PheRS are susceptible to reaction hijacking by DACM.

**Figure 3.**
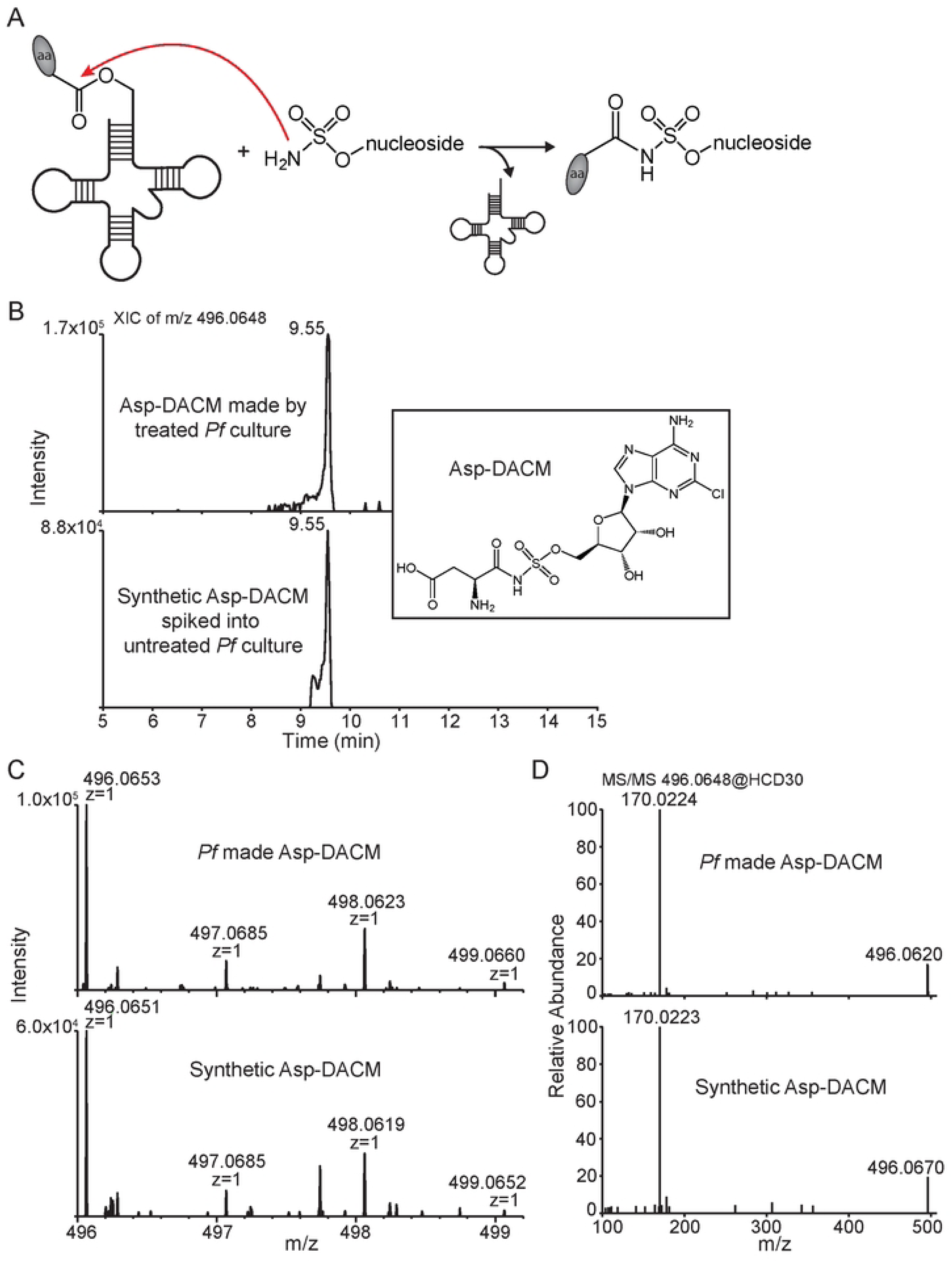
Targeted mass spectrometry identifies Asp-DACM conjugates in *P. falciparum*. (A) aaRSs catalyze nucleoside sulfamate attack on activated amino acids to form an amino acid adduct. (B-D). Late trophozoite stage *P. falciparum* 3D7 cultures were exposed to 10 μM DACM for 3 h. Parasite extracts were subjected to LC-MS/MS to search for DACM-amino acid conjugates. (B) Extracted ion chromatogram (XIC) of Asp-DACM *m/z* = 496.0648) from *P. falciparum* extracts and Asp-DACM standard spiked into untreated parasite lysate. Inset: Structure of Asp-DACM. (C,D) Mass spectra (C) and MS/MS fragmentation spectra (D) of Asp-DACM produced by *P. falciparum* extracts (top panels) and Asp-DACM standard (bottom panels).

### PfTyrRS is less susceptible to hijacking by DACM than by AMS

The low Tyr-DACM adduct signal suggested that *Pf*TyrRS was not a major target of DACM. This contrasts with AMS for which Tyr-AMS was the most easily detected product in mass spectrometry data [15]. We generated recombinant *Pf*TyrRS as described previously [17]. We used differential scanning fluorimetry (DSF) [17] to monitor changes in the thermal stability of recombinant *Pf*TyrRS upon incubation with AMS and DACM. Under the conditions examined, AMS exposure caused marked stabilization of *Pf*TyrRS (a 10°C shift), while DACM did not (Supp Fig. 3A; Supp Table 1), suggesting weaker hijacking activity. To understand why *Pf*TyrRS might be less susceptible to hijacking by DACM, we carried out protein–ligand docking using the Surflex fragment matching strategy [33]. Unconstrained docking always resulted in unrealistic orientations for Tyr-DACM, so the position of the Tyr fragment of Tyr-AMP-bound *Pf*TyrRS (PDB: 7ROR) was used to constrain the position of the Tyr-moiety in Tyr-AMP, Tyr-AMS and Tyr-DACM, which were docked into the active site *in silico* (without protein flexibility). Tyr-DACM gives the least favorable docking score (Supp Table S2), due to a clash of the Cl atom with surrounding residues (Supp Fig. 3B).

### Generation of recombinant AspRSs

*Pf*AspRS was chosen for further analysis as it has not been studied as a reaction-hijacking target previously. Alignment of the AspRS amino acid sequences from *P. falciparum*, *P. vivax, Saccharomyces cerevisiae* and humans reveals the characteristic anticodon-binding domain linked by a hinge region to the C-terminal catalytic domain, with conserved motifs 1 – 3 [34] (Fig. 4A; Supp Fig. 4). The *Plasmodium* sequences have a species-specific insertion in the anticodon-binding domain [35], while the *Plasmodium* and yeast sequences have a variable length N-terminal extension with a lysine-rich motif that is thought to bind RNA [35, 36]. Previous studies indicated that translation of *Pf*AspRS is initiated from an internal methionine, Met 49 [35].

**Figure 4.**
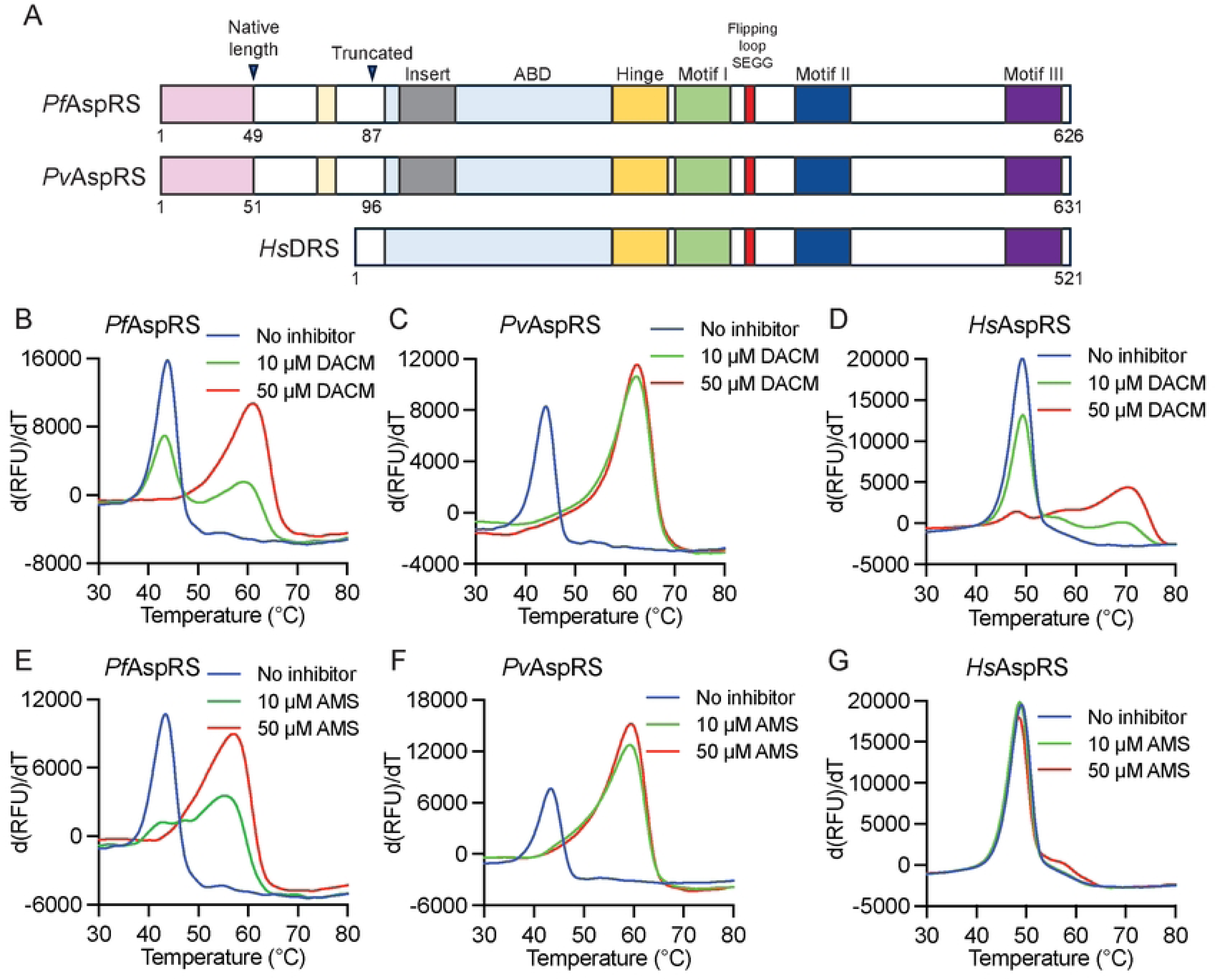
Thermal stabilization of native length AspRS enzymes by DACM and AMS. (A) Schematic diagram of AspRS enzymes from *P. falciparum, P. vivax* and *Homo sapiens*. Native length *Pf*AspRS and *Pv*AspRS are initiated from internal methionines, M49 and M51, respectively. The anticodon binding domain (ABD), the hinge region, Motifs I-III, and the flipping loop are indicated. Truncated *Pf*AspRS and *Pv*AspRS constructs comprise residues 87-626 and 96-631, respectively. (B-G) First derivatives of melting curves for native length *Pf*AspRS (B,E), *Pv*AspRS (C,F) and *Hs*AspRS (D,G) (1.5 μM) after incubation at 37°C for 3 h with 10 μM ATP, 20 μM Asp, 80 μM *Ec*tRNA, with 10 or 50 μM DACM (B-D) or 10 or 50 μM AMS (E-G). Data are representative of three independent experiments.

We used an *E. coli* expression system to generate recombinant native length *Pf*AspRS (49-626) (PF3D7_0102900) and the equivalent predicted native length *Pv*AspRS (51-631) (PVX_081610). Full-length *Hs*AspRS (1-501) (NP_001340.2) was also generated to enable a comparison of the susceptibility of the *Plasmodium* and human AspRS enzymes to reaction hijacking. Following removal of the His-tags, gel filtration yielded dimeric proteins. Given a previous report of poor stability of native length *Pf*AspRS [35], we characterized the preparation by mass spectrometry and analytical ultracentrifugation, showing that *Pf*AspRS exists as a homogenous dimer in solution (Supp Fig. 5A,B).

**Figure 5.**
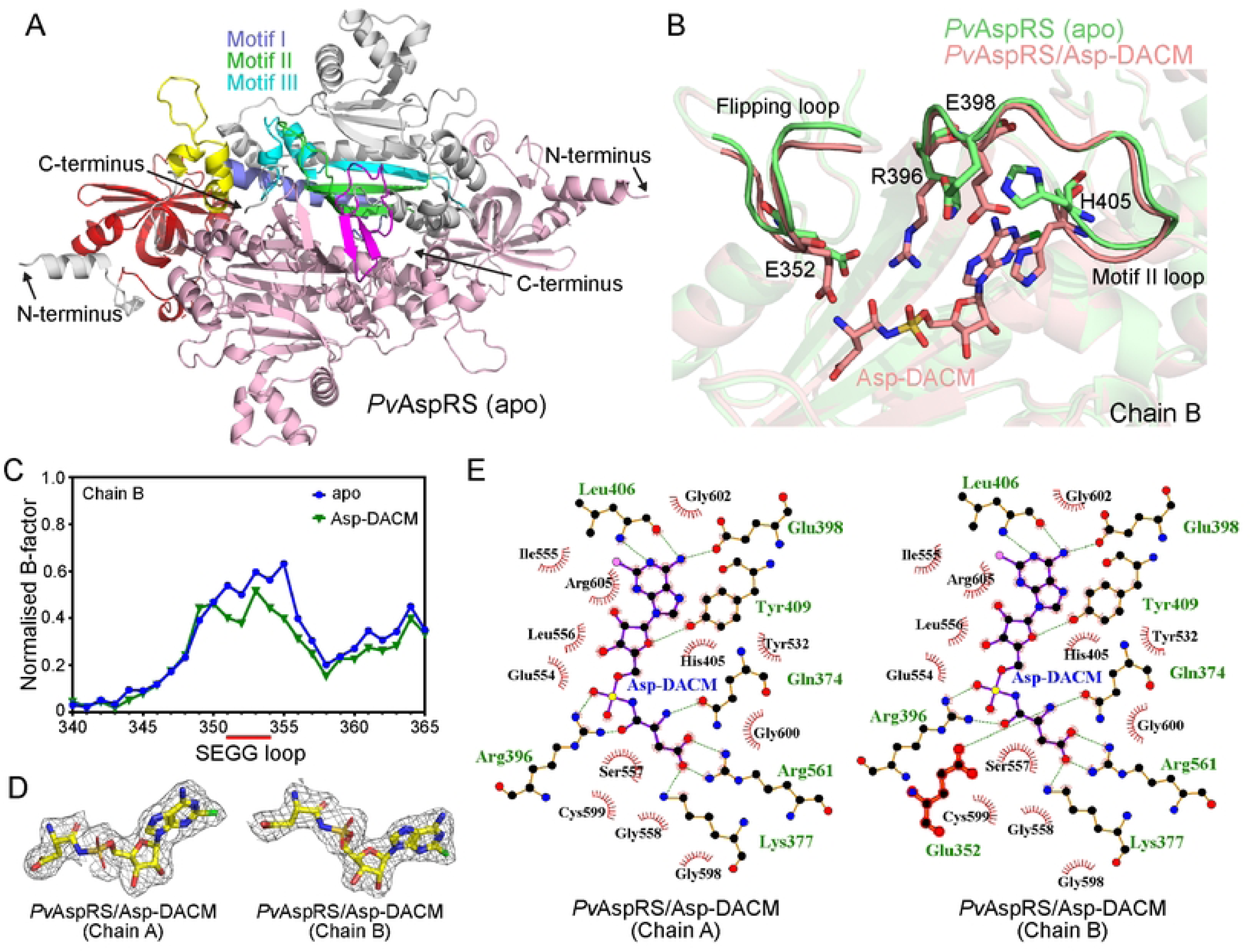
Comparison of the crystal structures of apo and Asp-DACM-bound *Pv*AspRS reveals binding site interactions. (A) Ribbon diagram of apo *Pv*AspRS (96-631) with Chain A in salmon and Chain B with domains indicated. The ABD (red), hinge region (yellow), motif I (violet), motif II (green), motif III (cyan) and flipping loop (magenta) are highlighted, with other regions in grey. (B) Overlay of ribbon representations of the flipping loop and the motif II loop of chain B of apo and Asp-DACM-bound *Pv*AspRS. The sidechain of residue R396 of apo *Pv*AspRS is not fully built due to insufficient density. (C) B-factor analysis of the B chain flipping loops of apo *Pv*AspRS and Asp-DACM-bound structures. The x-axis shows residue number. (D) 2*F*_o_-*F*_c_ maps contoured at 2 σ (mesh surface) showing electron density supporting the position of Asp-DACM bound to chains A and B. (E) Ligplots of Asp-DACM active site interfaces for the A and B chains. Hydrogen bonds and salt bridges are depicted with dashed (green) lines. Other interactions between protein and ligand are indicated by red arcs.

We assessed the ability of the recombinant aaRSs to consume ATP in the initial phase of the aminoacylation reaction, using the Kinase GLO assay, as previously described [17]. In initial studies, we found that laboratory prepared *E. coli* tRNA (*Ec*tRNA) is effective as a substrate for all three enzyme preparations, which facilitated the comparison. In the absence of tRNA, very little ATP is consumed by each of the enzymes (Supp Fig. 5C). Addition of *Ec*tRNA substantively increased the level of ATP consumption (Supp Fig. 5C), consistent with productive aminoacylation.

### Recombinant P. falciparum and P. vivax tRNA synthetases are thermally stabilized upon formation of Asp-DACM and Asp-AMS adducts

Upon incubation in the presence of substrates, *i.e.*, Asp, ATP and *Ec*tRNA, recombinant native length *Pf*AspRS, *Pv*AspRS and *Hs*AspRS exhibited melting temperature (*T*_m_) values of 43.0°C, 43.4°C, and 49.1°C, respectively (Fig. 4B-G; Supp Table 1). These *T*_m_ values are 2-3°C lower than for the respective apo AspRSs (Supp Table 1), which may be due to binding of the SYPRO Orange dye by the tRNA preparation, as reported previously [37].

When *Pf*AspRS was incubated with substrates in the presence of 50 μM DACM, the *T*_m_ value increased by 18.1°C (to 61.1°C), consistent with the formation of a very tightly bound Asp-DACM adduct (Fig. 4B; Supp Table 1). At a lower DACM concentration (10 μM), two peaks were evident indicating incomplete conversion to the stabilized form of *Pf*AspRS. *Pv*AspRS appears to be even more efficient at forming the Asp-DACM adduct, with the *T*_m_ value shift of 19.0°C (to 62.4°C) already largely complete with 10 μM DACM (Fig. 4C). By contrast, very little stabilization of *Hs*AspRS was observed at 10 μM DACM, but an emerging population of adduct-bound *Hs*AspRS was observed at 50 μM (Fig. 4D). As expected, incubation with synthetic Asp-DACM increased the *T*_m_ values of *Pf*AspRS, *Pv*AspRS and *Hs*AspRS to a similar extent (Supp Fig. 5D-F; Supp Table 1).

Incubation of the recombinant *Pf*AspRS and *Pv*AspRS enzymes with AMS plus substrates also led to marked stabilization, with increases in *T*_m_ values to 57.0 and 59.5°C, respectively. By contrast, *Hs*AspRS was not stabilized by AMS, under the conditions of this experiment (Fig. 4E-G; Supp Table 1). Taken together, these data suggest that *Hs*AspRS is less susceptible to reaction hijacking than *Pf*AspRS and *Pv*AspRS.

### Structure of apo PvAspRS

Attempts to crystallize native length *Pf*AspRS and *Pv*AspRS resulted in poorly diffracting crystals. An AlphaFold analysis predicted that the N-terminal extension was likely to be flexible (Supp Fig. 6A), which may impede crystallization. We therefore generated truncated *Pv*AspRS (96-631), lacking the N-terminal extension. *Pv*AspRS (96-631) appeared to be capable of catalyzing productive aminoacylation, as indicated by a marked increase in ATP consumption in the presence of *Ec*tRNA (Supp Fig. 6B). *Pv*AspRS (96-631) also appeared capable of generating the Asp-DACM adduct when incubated with 50 μM DACM in the presence of ATP, Asp and *Ec*tRNA, as indicated by an 18.4°C increase in the *T*_m_ value (Supp Fig. 6C, Supp Table 1). Incubation with the synthetic Asp-DACM adduct led to a 20.1°C increase in the *T*_m_ value (Supp Fig. 6D, Supp Table 1). These *T*_m_ values are similar to those for native length *Pv*AspRS, indicating that the core *Pv*AspRS construct binds the Asp-DACM adduct with similar affinity.

**Figure 6.**
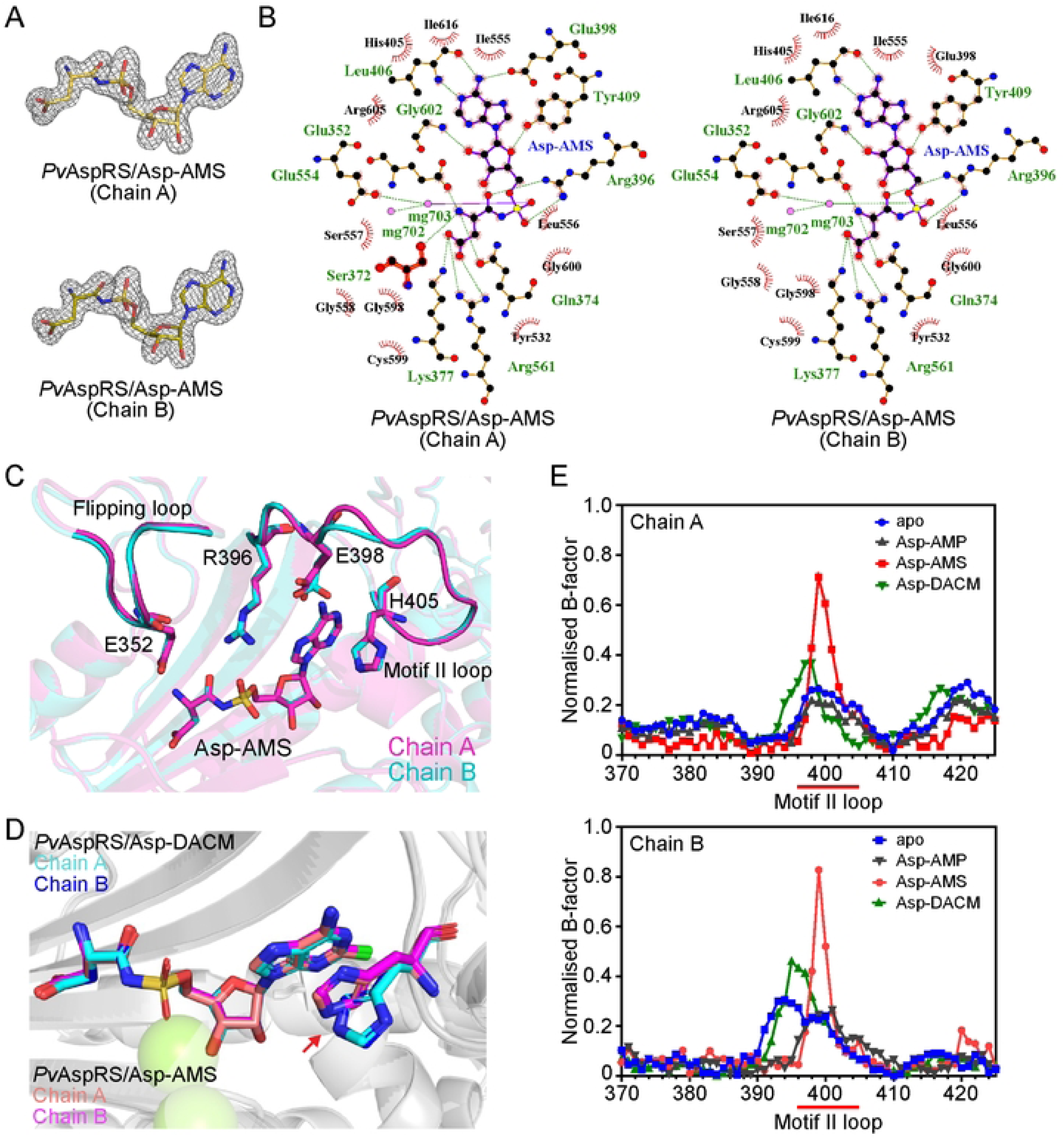
Crystal structure of Asp-AMS bound *Pv*AspRS. (A) 2*F*_o_-*F*_c_ maps contoured at 2 σ (mesh surface) showing electron density supporting the position of Asp-AMS bound to chain A and chain B. (B) Ligplots of Asp-AMS/active site interfaces for the A and B chains. Hydrogen bonds and salt bridges are depicted with dashed (green) lines. Other interactions between protein and ligand are indicated by red arcs. (C, D) Overlay of chains A and B of Asp-AMS-bound *Pv*AspRS showing the flipping loop and the motif II loop. (D) Overlay of chains A and B of Asp-AMS-bound and Asp-DACM-bound *Pv*AspRS, showing the different positions of H405. (E) B-factor analysis of the motif II loops of the apo and ligand-bound structures. The x-axis shows residue number.

A Morpheus II screen (Molecular Dimensions) was used to generate crystals of *Pv*AspRS (96-631), both as the apo protein and in complex with Asp-DACM, Asp-AMP and Asp-AMS. We solved the structure of apo *Pv*AspRS and refined it to 2.1 Å resolution. Data collection and refinement statistics are summarized in Supp Table 3. The structure of the apo dimer is presented in Fig. 5A, with features highlighted in Chain B, revealing a typical Type II aaRS with an N-terminal β-barrel anticodon-binding domain connected via a hinge to a larger C-terminal catalytic domain that adopts an α–β fold. Motif I is involved in the dimer interface. Motif II and motif III are integral components of the catalytic pocket.

The flipping loop, which plays an important role in substrate binding (Schmitt et al., 1998) [38], is supported by a β-hairpin structure (Fig. 5A, magenta). In apo *Pv*AspRS, residues SEGG (351-354) of the Chain A flipping loop are not resolved (Supp Fig. 7A). In Chain B, the flipping loop can be built (Fig. 5B). The SEGG loop interacts with a conserved residue Q371, as well as S350, N356 and A357 (Supp Fig. 7B). B-factor analysis suggests the SEGG loop is more mobile than neighboring regions of the protein (Fig. 5C).

**Figure 7.**
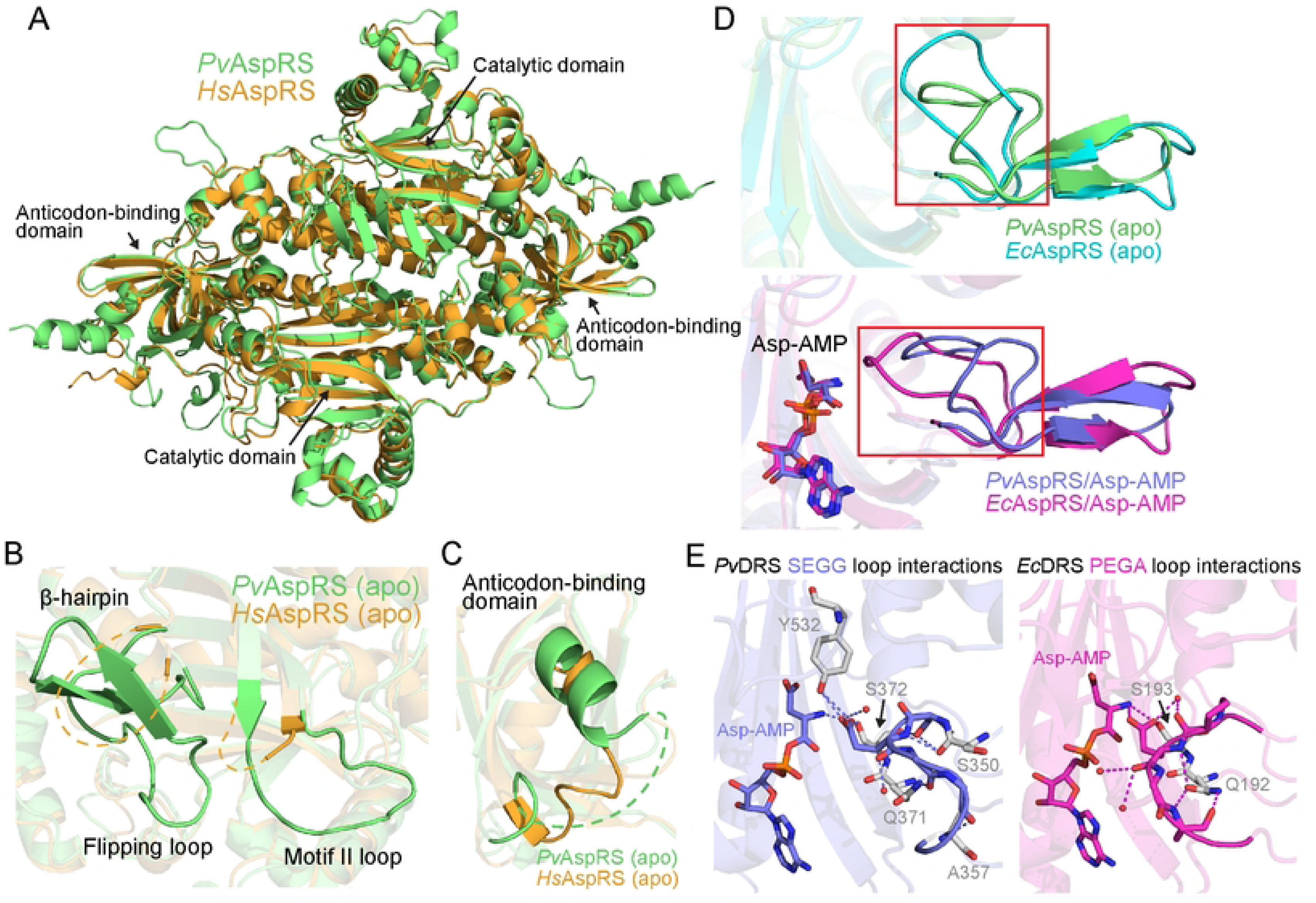
Comparison of the crystal structures of apo *Pv*AspRS with *Hs*AspRS and *Ec*AspRS. (A) Overlay of apo *Pv*AspRS and apo *Hs*AspRS (PDB: 4j15) illustrating the conservation of the overall structure. (B) The flipping loop and associated β-hairpin region, and the motif II loop, are unstructured in apo *Hs*AspRS and more ordered in apo *Pv*AspRS (Chain B). (C) Overlay of the anticodon binding domains of *Pv*AspRS and *Hs*AspRS. The *Plasmodium*-specific insertion is not resolved. (D) Overlay of the apo (top panel) and Asp-AMP-bound (bottom panel) *Pv*AspRS (chain B) and *Ec*AspRS (PDB: 1eqr chain A and 1il2 chain B) active sites illustrating the different positions of the flipping loops. (E) Comparison of the flipping loop interactions of Asp-AMP-bound *Pv*AspRS (SEGG; chain B) and *Ec*AspRS (PEGA; PDB: 1il2, chain B). SEGG interactions are depicted on the left panel. PEGA interactions are depicted in the right panel. SEGG and PEGA loop residues are shown as sticks and the interacting residues are shown in grey.

### Structure of Asp-DACM-bound PvAspRS

We solved the structure of *Pv*AspRS with bound synthetic Asp-DACM and refined it to 2.4 Å resolution (Supp Table 3). Well-defined electron density was observed for the adduct in both chains A and B (Fig. 5D). The Asp-DACM is located in the active site pocket of the catalytic domain and interacts with a number of amino acid residues (Fig. 5E). The adenine moiety is stacked between H405 and Y409 from motif II. H405 moves from its position in the apo protein to form a parallel π stack with the adenine ring (Fig 5B; Supp Fig. 7A). Other motif II loop residues, R396 and E398, also adopt different conformations in the Asp-DACM structure. R396 is highly conserved in prokaryotic and eukaryotic enzymes; and is known to stabilize the transition state during tRNA charging [39]. In both chains, R396 extends to interact with the aspartate of Asp-DACM (Fig. 5B,E; Supp Fig. 7A). Similarly, E398 repositions to form an H-bond with the adenine. The aspartic acid moiety of Asp-DACM also interacts with Q374, R561, and K377 (Fig. 5E).

As for apo *Pv*AspRS, the flipping loop in Chain B of Asp-DACM-bound *Pv*AspRS is well defined. Residue E352 (of the SEGG motif in Chain B) repositions to interact with the amino group of the aspartate, leading to a small movement towards the ligand (Fig. 5B). In Chain A of Asp-DACM-bound *Pv*AspRS, two additional residues of the flipping loop (G353 and G354) are resolved compared with the apo protein (Supp Fig. 7C).

### Structure of Asp-AMS-bound PvAspRS

We also solved the structure of *Pv*AspRS with bound synthetic Asp-AMS and refined it to 1.8 Å resolution, where well-defined electron density was observed for the adduct (Supp Table 3; Fig. 6A). Asp-AMS displays similar interactions to Asp-DACM (Fig. 6B), consistent with their similar ability to thermally stabilize *Pv*AspRS. Interestingly, in the Asp-AMS B chain structure, a Mg^2+^ ion coordinates the interaction between the sulfamate oxygen and E554 (Fig. 6B). E352 interacts with the amino group of the aspartate in both chains, leading to stabilization of both flipping loops (Fig. 6C). B-factor analysis reveals that the B chain flipping loop in Asp-AMS bound *Pv*AspRS exhibits higher stability than in the apo structure (Supp Fig 7C). Interestingly, H405 in the motif II loop adopts a different orientation to the Asp-DACM-bound structure (Fig. 6D). It forms a less intimate contact with the adenine ring that may be mediated by CH-π or cation-π interactions. Notably, B-factor analysis indicates that, in both chains, the motif II loop in Asp-AMS-bound *Pv*AspRS exhibits greater mobility than other structures (Fig. 6E). Thus, the interactions of Asp-DACM and Asp-AMS with the active site are subtly different.

### Structure of Asp-AMP-bound PvAspRS

We also solved the structure of *Pv*AspRS with the natural intermediate, Asp-AMP, and refined it to 2.1 Å resolution (Supp Table 3, Supp Fig. 8A). Asp-AMP is located in the equivalent position within the catalytic domain as Asp-DACM and Asp-AMS; and many of the interactions are similar (Supp Fig. 8B). Of interest, in the Asp-AMP structure, H405 occupies the same position as in the apo protein and does not form a π-π interaction with the ligand (Supp Fig. 8C). Similarly, E398 from the motif II loop remains in a similar position as in the apo protein, directed away from the active site (Supp Fig. 8C, right panel). This orientation is not consistent with E398 interacting with the amino group in the adenine ring, as is the case for the Asp-DACM and Asp-AMS structures (compare Fig. 6C and Supp Fig. 8C, right panel). As for apo *Pv*AspRS, the flipping loop SEGG residues are resolved in Chain B, but not in Chain A (Supp Fig. 8C). B-factor analysis shows that the B chain flipping loop in the Asp-AMP-bound structure has similar flexibility to the apo and Asp-DACM structures (Supp Fig. 7C); but is less ordered than the Asp-AMS structure (Supp Fig. 7C).

### Comparison of apo PvAspRS with apo HsAspRS

The structure of apo *Hs*AspRS (PDB: 4j15) has been published previously [40]. The two chains of the dimer are equivalent in the human structure. Superimposition of apo *Hs*AspRS with apo *Pv*AspRS (96-631) chain B reveals conservation of the overall structure, particularly in the catalytic and anticodon binding domains (Fig. 7A). By contrast, some regions of differential flexibility are observed (Fig. 7B,C). Of particular interest, residues I224-Q248, comprising the flipping loop and flanking residues, are disordered in *Hs*AspRS. By contrast, in the equivalent region of apo *Pv*AspRS (L347-Q371), the flipping loop and flanking β-hairpin are both well-defined in Chain B (Fig. 7B), while in apo Chain A, only the flipping loop itself is not resolved. Similarly, a loop (R273 to L282) in the middle of motif II, including residues that interact with the end of acceptor stem on the major groove [40–42], is not resolved in apo *Hs*AspRS, but the equivalent region (R396-H405) is ordered in apo *Pv*AspRS (Fig 7B). *Pv*AspRS exhibits an insert in the anticodon binding domain that is not present in *Hs*AspRS. The insert is disordered and not resolved in the *Pv*AspRS structure (Fig. 7C).

### Comparison of apo and Asp-AMP-bound structures of PvAspRS and EcAspRS

We also compared the apo and Asp-AMP-bound structures of *Pv*AspRS (96-631) with the equivalent *E. coli* AspRS structures. In apo *Ec*AspRS (PDB: 1eqr; [43]), the flipping loop is resolved, but with a high B factor. The apo *Ec*AspRS flipping loop (molecule 1/chain A) adopts an “open” conformation that is displaced compared with the flipping loop in chain B of *Pv*AspRS (Fig. 7D, top panel). By contrast, in the Asp-AMP-bound *Ec*AspRS structure (PDB: 1il2; [38]), the flipping loop adopts a “closed’’ structure (Fig. 7D, bottom panel) that is similar to that of Asp-AMP-bound *Pv*AspRS. In apo *Pv*AspRS, the SEGG loop is stabilized by interaction with the conserved residues Q371 and S372, as well as residues Y532, S350 and A357 (Fig. 7E). These latter residues are not conserved in *Ec*AspRS, and the equivalent PEGA loop only interacts with Q192, S193 and two water molecules. Thus, differences in the sequences of *Pv*AspRS and *Ec*AspRS in this region likely underpin the observed structural differences, enabling the apo *Pv*AspRS chain B flipping loop to adopt a conformation that is poised to form a “closed” structure.

## Discussion

Natural product nucleoside sulfamate antibiotics were identified nearly 70 years ago. Nucleocidin, a fluorinated nucleoside, was isolated from a soil dwelling *Streptomyces* [44, 45]. At the time, it was notable as the first compound, natural or synthetic, with an N-unsubstituted sulfamate ester group. Later, a chlorine-containing nucleoside sulfamate, DACM, was isolated from *Streptomyces* [19, 46]. These AMP mimics were shown to have activity against Gram-positive and Gram-negative bacteria and trypanosomes [18, 19, 44]. The synthetic AMP mimic, AMS, was also shown to be active against bacteria [22]. The mechanism of action of the natural nucleoside sulfamates was not clear, but they were known to inhibit protein synthesis [20–22, 47].

Here we show that DACM and AMS cause potent inhibition of the growth of *P. falciparum.* These compounds also show inhibitory activity against a mammalian cell line. This is in agreement with a previous study showing that DACM causes toxicity in mice [46]. Such toxicity will limit the development of these broad specificity nucleoside sulfamates as antimalarials. Nonetheless, AMS and DACM represent useful starting points for the synthesis of more selective compounds that can target the parasite AspRS.

Treatment of *P. falciparum* cultures with DACM or AMS inhibits protein translation and induces phosphorylation of eIF2α, as expected upon inhibition of aaRSs and the consequent accumulation of uncharged tRNA. EIK1, the *P. falciparum* homologue of GCN2, is the responsible kinase [48, 49]. Accordingly, treatment of EIK1 knockout parasites with DACM or AMS, did not result in eIF2α phosphorylation. Taken together, these data indicate that AMS and DACM exert their activity by inhibiting aaRSs.

Reaction hijacking of aaRSs represents a new and unusual mechanism to inhibit protein synthesis in *Plasmodium*-infected RBCs. Compounds that inhibit aaRSs via this mechanism induce the enzyme to synthesize amino acid adducts with a stable sulfamoyl bond. The adduct binds tightly in the active site and replaces the natural amino acid adenylate, which has a labile aminoacyl bond. Targeted mass spectrometry can be used to detect amino acid adducts of nucleoside sulfamates in treated *P. falciparum-*infected RBCs; and thus identify the susceptible target(s). In previous studies, we showed that ML901/ ML471 and OSM-S-106, specifically target *Pf*TyrRS (class I) and *Pf*AsnRS (class II), respectively [15–17]. By contrast, AMS targets several aaRSs, with *Pf*TyrRS being the most susceptible enzyme [15].

We interrogated the aaRS targets of DACM using mass spectrometry. The data indicate that *Pf*AsnRS, *Pf*AspRS, *Pf*ThrRS, *Pf*SerRS, *Pf*LysRS, *Pf*HisRS and *Pf*PheRS are susceptible to reaction hijacking by DACM. The apparently low activity against *Pf*TyrRS is interesting. Thermal stabilization studies confirmed that *Pf*TyrRS is less susceptible to hijacking by DACM than AMS. *In silico* analysis suggests that the 2-Cl substitution in DACM may clash with residues lining the adenylate pocket of *Pf*TyrRS, underpinning the observed selectivity.

Of the remaining targets of DACM, we considered *Plasmodium* AspRS to be worthy of additional study, as this enzyme has not been targeted for inhibitor development previously in *Plasmodium*; and no crystal structures were available before our study. *Plasmodium* AspRS is a Class IIb aaRS that exhibits unusual sequence features compared with human AspRS [35], including initiation from an internal methionine, and an insert in the anti-codon binding domain. *Plasmodium* AspRS also exhibits an extended N-terminal region, with a lysine-rich motif that is thought to facilitate tRNA binding [35]. *Hs*AspRS has a much shorter (21-residue) N-terminal extension that may also be involved in tRNA binding [50, 51].

We generated recombinant, native length *Pf*AspRS (49-626) and the equivalent predicted native-length *Pv*AspRS (51-631), as well as a truncated construct of *Pv*AspRS (96-631), lacking the N-terminal extension. A previous study reported that native length *Pf*AspRS exhibits a propensity to aggregate [35]. However, in our hands, this construct was well behaved in solution, as assessed by analytical centrifugation. We also generated *Hs*AspRS for comparison. Each of these constructs appear to be capable of catalyzing aminoacylation of tRNA, as judged by a substantive increase in ATP consumption upon addition of *Ec*tRNA. These data suggest that the N-terminal extension is not essential for enzyme activity.

Upon incubation of the recombinant AspRS constructs with DACM or AMS in the presence of all substrates, a substantial increase in the apparent melting temperatures is observed, consistent with the formation of very tightly bound Asp-DACM and Asp-AMS adducts via the reaction hijacking mechanism. By contrast, *Hs*AspRS appears to be less susceptible to hijacking, particularly by AMS. This is consistent with previous studies showing that *Pf*TyrRS and *Pf*AsnRS are more susceptible to hijacking than their human counterparts [15–17].

We solved the first structures of AspRS from *Plasmodium. Pv*AspRS (96-631) was successfully crystallized as the apo protein, in complex with the natural intermediate, and with synthetic Asp-AMS and Asp-DACM, allowing insights into the binding determinants. As anticipated, *Pv*AspRS adopts a dimeric structure with features typical of Type II aaRSs, namely an N-terminal anticodon-binding domain connected via a hinge domain to a C-terminal catalytic domain, with characteristic motifs (I-III) and a flipping loop over the active site. The *Plasmodium*-specific insertion in the ABD (residues 137 – 167) is located at the protein surface and resides 159-179 are unresolved. This region is distal to the predicted tRNA anticodon binding surface and the role of this insertion remains unclear.

AspRSs have been studied extensively in bacteria, yeast and humans. The active site comprises binding pockets for Asp, AMP and the 3ʹ end of tRNA. In apo *Hs*AspRS and *Pyrococcus kodakaraensis* AspRS, the flipping loop and flanking residues are disordered [40, 52]. In the apo forms of the yeast and *E. coli* AspRS, the flipping loop adopts a defined open conformation [43, 53]. In each of these cases, following binding of the amino acid substrate, the flipping loop adopts a closed lid-like conformation that contributes to positioning the amino acid in a state ready for attack [38, 52]. The closed loop also prevents access of the terminal tRNA adenosine until the adenylate intermediate is formed. The flipping loop then repositions to allow access of the tRNA 3ʹ end; and it then anchors the A76 base to promote the transfer step [38].

In contrast to many other species studied to date, *Pv*AspRS exhibits structural asymmetry even in the apo state. In *Pv*AspRS chain A, the flipping loop is disordered, while in chain B, the density could be mapped, although it exhibits higher B-factor values, indicating some flexibility of the loop. Interestingly, only minor shifts are observed after binding of ligands suggesting that the apo *Pv*AspRS chain B flipping loop is poised to adopt a closed position. Upon binding of Asp-AMP, Asp-DACM or Asp-AMS, E352 (of SEGG) undergoes modest repositioning to interact with the amino group of the aspartate. In chain A of Asp-AMS-bound *Pv*AspRS, the flipping loop becomes structured and E352 engages with the ligand, while in Asp-DACM *Pv*AspRS chain A, only the GG of the flipping loop SEGG residues are resolved.

Upon binding of either Asp-DACM or Asp-AMS, changes are also observed in the motif II loop. E398 extends towards the adenine and forms an interaction with the amino group. In contrast, in Asp-AMP-bound *Pv*AspRS, E398 remains in a similar position as in the apo protein, directed away from the active site. This difference may contribute to the tighter binding of the nucleosulfamide adducts. The tighter binding of Asp-DACM or Asp-AMS is also likely enhanced by the stable sulfamoyl bond of the nucleosulfamide adducts, which prevents cleavage and effectively locks *Pv*AspRS in the adduct-bound state. Of interest, H405 adopts different orientations in the Asp-DACM or Asp-AMS structures, forming, respectively, either a parallel π-stack or a T-shaped interaction, that may be CH-π or cation-π in nature. These differences are associated with different flexibility of the motif II loop. The chloro group of DACM is expected to make the adenine ring more electron deficient, which will enhance the π-π interaction between H405 and the nucleobase.

In apo *Hs*AspRS, the motif II loop (R273 to L282) is not resolved. This loop includes residues that interact with the 3’ end of the tRNA acceptor stem [41, 42, 53]. By contrast, in apo *Pv*AspRS, the equivalent region (R396-H405) is ordered. Enhanced binding of aminoacylated tRNA could increase the longevity of the product-bound form of the enzyme, which could predispose *Pv*AspRS to attack by nucleoside sulfamates [15].

In conclusion, we have identified DACM as a new reaction hijacking inhibitor with potent activity against *P. falciparum.* The *Plasmodium* AspRS is revealed here as a new target for antimalarials that is more susceptible to reaction hijacking than its human counterpart. Differences in the conformation and flexibility of the flipping loop and the motif II loop may underpin the differential susceptibility between parasite and host enzymes. Further work is needed to develop more selective inhibitors that could be developed as future antimalarial compounds.

## Methods

### Inhibition of the growth P. falciparum cultures

Sorbitol-synchronized ring stage parasites at 1% parasitaemia and 0.2% haematocrit (3D7 strain [54]) were incubated with DACM or AMS for 72 h and viability was assessed in the next cycle by flow cytometry, after labelling with 2 μM Syto-61 (Thermo Fisher Scientific), as previously described [28]. The parasitemia was normalized to untreated and “kill-treated” controls, treated with 10x IC_50_ concentration of each compound, for 72 h. Cells were pelleted at 400 *g* for 1.5 min and were incubated with 2 μM SYTO 61 in phosphate-buffered saline (PBS; Molecular Probes, Life Technologies) at RT for 15 min. Nine volumes of PBS were added to the cells (final SYTO 61 concentration: 0.2 μM) and incubated for a further 30 min at RT. The samples were analysed by BD Biosciences FACSCanto™ II flow cytometer using the APC channel where the forward and side scatter was used to gate total cells. Data were processed by BD FACSDiva Software and FlowJo™. The half maximal inhibitory concentration (IC_50_) was determined using nonlinear regression (curve fit) in GraphPad Prism.

### Cytotoxicity of PM03 against HepG2 cells

The HepG2 (Human Caucasian hepatocyte carcinoma) cell line was procured from Cell Repository NCCS, India, and cultured in Dulbecco’s Modified Eagle Medium (DMEM) supplemented with 10% Fetal Bovine Serum (FBS), 4 mM glutamine and 50 μg/mL penicillin-streptomycin, in a humidified incubator at 37°C with 5% CO_2._ Using an assay procedure modified from [55], 10,000 cells per well were incubated in 96-well plates for 24 h at 37°C with 5% CO_2_, to allow attachment. The medium was removed; and cells were treated with fresh medium (100 μL) containing either vehicle or serial dilutions (2-fold) of the compound, prepared in DMEM, with 2% FBS, in triplicate. Cells treated with 20% DMSO were used as a kill control. The cells were incubated for 48 h at 37°C with 5% CO_2_. The growth medium was aspirated, and 100 μL of 0.5 mg/mL MTT solution was added to each well. After 3 h at 37°C, with 5% CO_2_, the MTT solution was removed and 100% DMSO was added to dissolve formazan crystals, with shaking, in the dark, at 37°C for 15 min. The absorbance was measured at 570 nm using a SpectraMax M3 microplate reader. The absorbance at 630 nm was subtracted to correct for background noise. Graph Prism 9 was used to generate the dose-response curve using non-linear regression analysis (variable slope).

### Protein translation assay

*P. falciparum* Cam 3.II_Rev [56] trophozoite (30-35 h p.i.) infected RBCs (0.2% hematocrit and 1% parasitemia) were exposed to the relevant compounds for 6 h. The cells were labelled with O-propargyl-puromycin (OPP (Abcam); 4 μM) in the final 2 h, washed two times (PBS + 3% human serum) and then fixed and permeabilized, as described previously [17]. The click reaction was performed for 1 h at 37°C in the presence of CuSO_4_ (0.1 mM), tris-hydroxypropyltriazolylmethylamine (THPTA; 0.5 mM) and sodium ascorbate (5 mM) to bring about azide-alkyne cycloaddition to Alexa Fluor 488 azide (0.1 μM). Pellets were washed four times and resuspended in PBS + 3% human serum containing 25 μg/ml propidium iodide (Invitrogen™). Flow cytometry (FACS Canto II; BD Biosciences, San Jose, CA) was performed using a BD FACSDiva (version 8.0) and FlowJo (version 10.9). Side scatter height (SSC-H) and forward scatter area (FSC-A) density plots were used to gate the total cell population. FSC-A and forward scatter width (FSC-W) plots were used to gate the single cell population. The FITC and PerCP-Cy5.5-H channels were used to detect the Alexa Fluor 488 and propidium iodide (PI) positive populations, representing parasitized red blood cells. The same 6-h drug treatments conditions were set up in parallel for viability assessment. Cells were washed intensively 3 x with Complete Culture Medium (CCM) to remove inhibitors and returned to culture. Parasite viability was assessed in the next cycle as previously described above.

### Phospho-eIF2α analysis

Trophozoite-infected RBCs (26 – 32 h p.i.; 2.5% hematocrit, 5-6% parasitemia) of the Cam 3.II_Rev line [56] or an *eIK1* knock-out line [29] (kindly provided by Prof Christian Doerig, RMIT University), were incubated with 0.2 μM borrelidin, 1 μM ML901, 1 μM AMS, 1 μM DACM, or DMSO, for 3 h. The RBC pellet was washed 3 x in PBS + cOmplete EDTA-free Protease Inhibitor Cocktail. The pelleted cells were lysed by resuspension in PBS + 0.05% saponin, on ice. Washed pellets were resuspended in Bolt LDS sample buffer plus reducing agent and subjected to Western analysis as described previously [17]. Primary antibodies: rabbit anti-phospho-eIF2α (Cell Signaling Technology-119A11; 1:1,000); polyclonal mouse anti-*Pf*BiP, generated using recombinant *Pf*BiP at the WEHI Antibody Services (1:1,000). Secondary antibodies: goat anti-rabbit IgG-peroxidase (Chemicon-A132P; 1:20,000), goat anti-mouse IgG-peroxidase (Chemicon-A181P; 1:50,000). Membranes were washed three times in 1xPBS + 0.1% Tween 20. Chemiluminescence was detected using the Bio-Rad ChemiDoc^TM^ MP imaging system.

### Sample preparation to identify amino acid DACM conjugates

Late trophozoite stage *P. falciparum* (3D7 strain) culture samples was exposed to 10 μM DACM for 3 h. The parasite-infected RBCs were lysed with 0.1% saponin in PBS and the parasite pellet was washed 3 times with ice-cold PBS. Cell pellets were kept on ice and resuspended in water as one volume, followed by the addition of five volumes of cold chloroform-methanol (2:1 [vol/vol]) solution. Samples were incubated on ice for 5 min, subjected to vortex mixing for 1 min and centrifuged at 13,500 x *g* for 10 min at 4°C to form 2 phases. The top aqueous layer was transferred to a new tube and subjected to LC-MS analysis. The synthetic Asp-DACM standard was processed in the same way. Data analysis was performed using Xcalibur (version 4.4).

### High-performance liquid chromatography (HPLC) and mass spectrometric (MS) analyses

Samples were analysed by reversed-phase ultra-high performance liquid chromatography (UHPLC) coupled to tandem mass spectrometry (MS/MS) employing a Vanquish UHPLC linked to an Orbitrap Fusion Lumos mass spectrometer (Thermo Fisher Scientific, San Jose, CA, USA) operated in positive ion mode, modified from a previous procedure [15]. Solvent A was 0.1% formic acid acetate in water and solvent B was 0.1% formic acid in acetonitrile. 5 μL of each sample was injected onto an RRHD Eclipse Plus C18 column (2.1 × 100 mm, 1.8 μm; Agilent Technologies, USA) held at 50 °C with a solvent flow rate of 350 μL/min. The solvent gradient was as follows [Time (min), B %]: [0, 0], [6, 0], [13, 25], [13.1, 99], [14, 99], [14.1, 0], [18, 0]. Mass Spectrometry experiments were performed using a Heated Electrospray Ionization (HESI) source. The spray voltage, flow rate of sheath, auxiliary and sweep gases were 3.5 kV, 25, 5, and 0 ‘arbitrary’ unit(s), respectively. The ion transfer tube and vaporizer temperatures were maintained at 300°C and 150°C, respectively, and the S-Lens RF level was set at 30%. A full-scan MS spectrum and targeted MS/MS for the proton adduct of Asp-DACM or the 20 possible common amino acid-containing inhibitor adducts were acquired in cycles throughout the run. The full-scan MS-spectra were acquired in the Orbitrap at a mass resolving power of 120,000 (at *m/z* 200) across an *m/z* range of 200–1500 using quadrupole isolation and the targeted MS/MS were acquired using higher-energy collisional dissociation (HCD)-MS/MS in the Orbitrap at a mass resolving power of 7500 (at *m/z* 200), a stepped normalized collision energy (NCE) of 15, 30 and 45% and an *m/z* isolation window of 1.2.

### Generation of recombinant aaRSs

The gene sequences encoding native length *Pf*AspRS (49-626) (PlasmoDB ID: PF3D7_0102900), native length *Pv*AspRS (51-631) (PVX_081610) and full-length *Hs*AspRS (1-501) (NP_001340.2) were codon-optimized for expression in *Escherichia coli*, synthesized by GenScript, and cloned into pET-11a vector with a histidine tag and TEV cleavage site. Truncated *Pf*AspRS (87-626) and *Pv*AspRS (96-631) genes were amplified from the synthesized plasmids using PCR and cloned into pET-11a and pETM-41 vectors, respectively. The proteins were overexpressed in *E. coli* BL21 (DE3) using 0.05 mM or 0.1 mM IPTG induction at 16°C, overnight. The lysis buffer was 50 mM Tris pH 7.4 (or pH 8.0), 350 mM NaCl, 40 mM imidazole, 0.5 mM tris(2-carboxyethyl)phosphine (TCEP), 1 mg/mL lysozyme, and 1x complete protease inhibitor. Clarified lysate was loaded onto a HisTrap (HP) 5 mL nickel column (Cytiva). Proteins were eluted with a gradient of Buffer B (50 mM Tris pH 7.4, 350 mM NaCl, 500 mM imidazole, 0.5 mM TCEP). *Pf*AspRS was eluted at ∼40% Buffer B. The eluted sample was dialyzed against 50 mM Tris-HCl pH 7.4, 150 mM NaCl, 25 mM imidazole (or 200 mM imidazole), and 1 mM TCEP, with TEV protease to remove the N-terminal histidine tag. The sample was concentrated and subjected to size exclusion chromatography using buffer containing 25 mM or 50 mM Tris-HCl pH 8.0, 150 mM NaCl, and 0.5 or 1 mM TCEP.

### Characterization of native length PfAspRS

For analytical ultracentrifugation, recombinant *Pf*AspRS was prepared at 1, 0.6, and 0.2 mg/mL in buffer containing 25 mM Tris pH 8.0, 150 mM NaCl, 1 mM TCEP. Samples and buffer (reference solution) were centrifuged at 200,000 g at 20°C, in a Beckman Coulter XL-I analytical ultracentrifuge, equipped with UV-visible scanning optics. Radial absorbance data were monitored and collected at a wavelength of 290 nm. Sedimentation data were fitted to a continuous sedimentation coefficient (c(s)) model, with frictional ratios estimated using SEDFIT software [57].

For mass spectrometry, purified protein (5 μg) in 25 mM Tris pH 8.0, 150 mM NaCl, 1 mM TCEP was subjected to an Agilent 1200 HPLC, equipped with a C18 column, connected to an Agilent 6220 Accurate-Mass electrospray ionization time-of-flight (ESI-TOF) mass spectrometer. Data acquisition and analysis were performed using Mass Hunter Software (Agilent).

### Preparation of E. coli tRNA

Total tRNA from *E. coli* was isolated with modifications from a previous report [58]. *E. coli* BL21(DE3) cells were cultured in 2× Yeast Extract Tryptone medium at 37℃ overnight. Cells were harvested by centrifugation, resuspended in diethyl pyrocarbonate (DEPC)-treated water, and lysed by adding TRIzol™ reagent (Invitrogen™) at a 3:1 ratio to the cell suspension, followed by vigorous vortexing. After centrifugation, the aqueous phase was extracted, and acid-phenol: chloroform pH 4.5 (with indole-3-acetic acid (IAA), 25:24:21; Invitrogen™) was added. Following another centrifugation step, the supernatant was collected. This step was repeated until a clear interface was observed. Next, LiCl was added to a final concentration of 1 M. tRNA was precipitated with ice-cold isopropanol, then dissolved in DEPC-treated water for further use.

### ATP consumption assay

The consumption of ATP by native length *Pf*AspRS, native length *Pv*AspRS, truncated *Pv*AspRS and full-length *Hs*AspRS was determined using a luciferase-based assay as per the manufacturer’s instructions (Kinase-Glo Luminescent Kinase Assay, Promega). Reactions were conducted in 25 mM Tris-HCl (pH 8), 150 mM NaCl, 5 mM MgCl_2_, 0.1 mg/mL BSA, 1 mM TCEP, with 200 μM L-aspartate, 10 μM ATP, 1 unit/mL inorganic pyrophosphatase and 80 or 160 μM *E.coli* tRNA (if present). Enzyme concentration and incubation time for each experiment are described in the figure legends. Reactions were incubated at 37°C, followed by addition of the Kinase-Glo reagent. Luminescence output was measured using a plate reader (CLARIOstar, BMG LABTECH) and the highest signal within 20 min after addition of reagents was recorded using MARS data analysis software (version 3.32). The concentration of ATP was quantified by linear regression using an ATP standard curve (Microsoft Excel).

### Differential scanning fluorimetry (DSF)

The effect of DACM, AMS and Asp-DACM on the thermal stability of AspRS enzymes was assayed as previously described [15]. Briefly, the relevant AspRS (1.5 μM) was incubated in the presence or absence of 10-50 μM DACM or AMS, or 2, 5 or 10 μM Asp-DACM, with 10 μM ATP, 20 μM L-aspartate, 80 μM *Ec*tRNA, in 25 mM Tris-HCl (pH 8), 150 mM NaCl, 5 mM MgCl_2_, 1 mM TCEP, at 37°C for 3 h. SYPRO Orange (Sigma-Aldrich; 5,000X concentrate in DMSO) was added to the reaction mixture at a final concentration of 5X. 25 μL of the sample was added into each well of a 96-well qPCR plate (Applied Biosystems). The plate was sealed and analysed using StepOnePlus Real-Time PCR system (Applied Biosystems). The samples were heated from 20°C to 90°C with a 1°C per min continuous gradient. The thermal unfolding curve was plotted as the first derivative curve of the raw fluorescence values. The melting temperature (*T*_m_), defined as the peak of the first derivative curve, was used to assess the thermal stability of protein-ligand complexes.

### Crystallization and X-ray diffraction data collection

For crystallization, *Pv*AspRS (96-631) in the apo form, prepared in 20 mM HEPES pH 8.0, 200 mM NaCl, 10% glycerol, and 5 mM beta-mercaptoethanol, was concentrated to 20 mg/mL and incubated without or with bound natural substrates (ATP and L-Asp) or synthetic Asp-AMS or Asp-DACM, with 5 mM MgCl_2_, at a molar ratio of 1:4 to 1:30 (*Pv*AspRS monomer/ ligand). Crystallization experiments were performed using the sitting-drop or hanging-drop vapour-diffusion method at 293 K. Initial crystallization screening was carried out in a 96-well plate (Corning, Lowell, Massachusetts, USA) with ViewDrop II seals (SPT LabTech, Melbourn, England) using the commercially available crystallization sparse-matrix screens (SG1, ProPlex, and Morpheus I and II; Molecular Dimensions, UK) [59]. Three different drop ratios were aliquoted using a Mosquito nanolitre dispenser system (TTP LabTech, Melbourn, England) or an NT8® drop setter (Formulatrix) by mixing protein and reservoir solutions at 1:1, 2:1 and 1:2 drop ratios; final volume 150 nL). The crystallization droplets were equilibrated against a 75 µL reservoir solution. Initial crystals of *Pv*AspRS were obtained in the following conditions. Apo *Pv*AspRS: SG1-D10 (0.2 M lithium sulfate, 0.1 M Bis-Tris pH 6.5 and 25% w/v PEG 3350); *Pv*AspRS (Asp-AMP): SG1-C12 (0.2 M sodium acetate trihydrate, 0.1 M Bis-Tris pH 5.5 and 25% w/v PEG 3350); *Pv*AspRS (Asp-AMS): Morpheus I-A9 (0.06 M divalents (0.3 M magnesium chloride hexahydrate; 0.3 M calcium chloride dihydrate), 0.1 M Buffer System (Tris base and BICINE), pH 8.5 and 30% v/v Precipitant Mix (40% v/v PEG 500 MME and 20% w/v PEG 20000); *Pv*AspRS (Asp-DACM): Morpheus II-B5 (15% (w/v) PEG 3000, 20% (v/v) 1,2,4-butanetriol, 1% (w/v) nondetergent sulfobetaine (NDSB) 256, 0.5 mM manganese chloride, 0.5 mM cobalt chloride, and 0.5 mM zinc chloride). The crystals were cryoprotected using 10-20% glycerol before being flash-cooled in liquid nitrogen. X-ray diffraction experiments were conducted on the MX2 beamline at the Australian Synchrotron [60] or the I03 beamline at the Diamond Light Source, UK (Supp Table 3).

### Structure determination

Several datasets were collected. Data were indexed and integrated using the XDS software package [61] and scaled using AIMLESS [62]. Alternatively, data were processed using the xia2/DIALS [63] and autoPROC [64] pipeline. Human AspRS (*Hs*AspRS, PDB ID: 4J15 [40]) was used as the phasing model. The initial phases were determined by molecular replacement using *PHASER* [65] or Auto-Rickshaw [66]. The model was further refined using *phenix.refine* from *PHENIX* [67, 68] and manually built using *COOT* [69]. Ligands, ions and water molecules were added to their electron densities after several rounds of manual model building and refinement. Structure refinement was performed using non-crystallographic torsion restraints and translation/libration/screw (TLS) refinement with each chain comprising a single TLS group. Restraints for Asp-AMP, Asp-AMS, Asp-DACM were generated using phenix.elbow [70] or GRADE Web Server (Global Phasing Ltd, https://grade.globalphasing.org). Difference density peaks observed near the 2-Cl moiety of Asp-DACM suggested radiation damage in this location during data collection. *MolProbity* [71, 72] in *PHENIX* suite was used to evaluate model quality and figures were generated using UCSF Chimera [73] and PyMOL (http://www.pymol.org), including the embedded PyMOL secondary structure assignment. Magnesium ions were identified using CheckMyMetal [74]. Complete data collection and refinement statistics are summarized in Supp Table 3. LigPlot+ (version 2.2.8) was employed to analyze ligand-protein interactions and to generate 2D graphical maps [75].

### Isotropic B-factor analysis

Isotropic B-factors for the alpha carbons for each residue were extracted from the PDB files in PyMOL Version 2.5.4 [75]. The B-factors were corrected by dividing by the Wilson B-factor of each structure 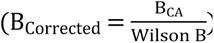 and then normalised using the following equation: 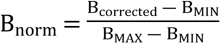. The resulting normalized, dimensionless B-factor derived values (B_norm_) ranged from 0-1, with higher values indicating greater atomic mobility.

### Chemistry

Synthetic procedures and compound characterizations are provided in the Supplementary Information.

## Acknowledgements

AMS was kindly provided by Steven Langston, Drug Discovery Sciences, Takeda Pharmaceuticals International Company, Cambridge, United States. We would like to thank the University of Melbourne Bio21 Molecular Science and Biotechnology Institute’s Melbourne Mass Spectrometry and Proteomics Facility, the Bio21-WEHI crystallization facility, and the Melbourne Protein Characterization Facility, including Roxanne Smith and Yee-Foong Mok, for technical support. This research was partly undertaken at the Australian Synchrotron, part of the Australian Nuclear Science and Technology Organization, and made use of the ACRF Detector on the MX2 beamline. We thank the beamline staff for their assistance. We are grateful for beamtime at the Diamond Light Source (DLS) and the staff of beamline I03 for data collection (BAG application mx28534). We are also grateful for beamtime at the SOLEIL beamlines PROXIMA 1 and 2A, and the staff of beamlines for their aid in preliminary data collection.

## Data Availability

All relevant data are within the manuscript and its Supporting Information files. The following structures have been deposited in the PDB: *Pv*AspRS-apo (9M5M); *Pv*AspRS:Asp-AMP (9M5N), *Pv*AspRS:Asp-AMS (9M5O); *Pv*AspRS:Asp-DACM (9NPJ).

## Author Contributions

Conceptualization: N.K., C-W.T., V.K.S, Y.M., Y.K., C.D., S-A. P., M.D.W.G., S.P., A.S., L.T., S.C.X; Investigation: N.K., C-W.T., V.K.S, Y.M., Y.K., W.T., C.D., D.H.H., C.J.M., A-S.C.B., M.L., B.S., P.I.P., M.D.W.G, S.C.X.; Analysis: N.K., C-W.T., V.K.S, Y.M., Y.K., W.T., C.D., D.H.H., C.J.M., S.P., B.S., P.I.P., S-A.P., M.D.W.G., A.S., L.T. S.C.X.; Funding acquisition: S-A. P., M.D.W.G., A.S., L.T., S.C.X.; Writing: N.K., C-W.T., V.K.S, Y.M., Y.K., D.H.H., S.P., B.S., P.I.P., S-A.P., M.D.W.G., A.S., L.T., S.C.X.

## Supplementary Figure Legends

Supplementary Figure 1. Activity of AMS against *P. falciparum* and HepG2 cells.

(A) Structure of AMS. (B) Sorbitol-synchronized ring stage parasites were subjected to a 72-h drug sensitivity assay with AMS (white circles). Data represent five independent experiments, each performed in duplicate. Cytotoxicity of AMS (black circles) against the HepG2 mammalian cell line. Data represent five independent experiments, each performed in triplicate. Error bars indicate SEM.

Supplementary Figure 2. Identification of amino acid-DACM-conjugates in *P. falciparum*.

(A) *P. falciparum* cultures were exposed to 10 μM DACM for 3 h. Parasite extracts were subjected to LC-MS/MS to search for DACM-amino acid conjugates. (A-F) Detected (top panels) and predicted mass spectra of (A) DACM-Asn (*m/z* = 495.0808), (B) DACM-Lys (*m/z* = 509.1328); (C) DACM-Thr (*m/z* = 481.0855); (D) DACM-Ser (*m/z* = 468.0699), (E) DACM-His (*m/z* = 518.0968); (F) DACM-Phe (*m/z* = 528.1063). (G-L). MS/MS spectra of the fragmented ions, including a *m/z* of 170.0228 found in fragmented ion of each DACM-amino acid conjugate.

Supplementary Figure 3. DSF and docking analysis for *Pf*TyrRS.

(A) First derivatives of melting curves of *Pf*TyrRS (2.3 μM) after incubation at 37°C for 2 h with 10 μM ATP, 20 μM Tyr, 4 μM *Pf*tRNA^Tyr^, with 5 or 10 μM of AMS or DACM. Data are representative of three independent experiments. (B) A chlorine atom was added to the 2-position of the adenine ring system of Tyr-AMP at the active site of *Pf*TyrRS (PDB: 7ROR) using ChimeraX [77]. Steric overlaps between the chlorine atom and the protein binding pocket are shown as red dashed rods.

Supplementary Figure 4. Sequence alignment of AspRS sequences from different species. Alignment of AspRS sequences from *P. falciparum* (*Pf*), *P. vivax* (*Pv*), *Homo sapiens* (*Hs*), *Saccharomyces cerevisiae* (*Sc*), reveals a high level of conservation of the three Type II aaRS motifs (I-III, blue, green, purple text), which are involved in ATP binding and dimerization. The hinge region is highlighted in yellow. The *Plasmodium* sequences exhibit a large N-terminal extension with native initiation from an internal methionine (aqua text). The anticodon-binding domain (salmon) has a *Plasmodium*-specific insert (underlined). The flipping loop residues, SEGG, that have previously been shown to undergo dynamic motions that facilitate tRNA binding [52], are in red text. Two loops that are ordered in *Pv*AspRS but disordered in *Hs*AspRS are boxed, namely the flipping loop and flanking β-hairpin structure and the motif II loop.

Supplementary Figure 5. Characterization of native length *Pf*AspRS and stabilization by the Asp-DACM adduct.

(A) Deconvoluted mass spectrum obtained using Agilent Mass Hunter software. The mass of the highest peak (67831 Da) correlates well with the theoretical mass of native length *Pf*AspRS. (B) Sedimentation velocity analysis. Purified native length *Pf*AspRS was diluted to 1, 0.6, and 0.2 mg/mL in buffer containing 50 mM Tris-HCl pH8.0, 150 mM NaCl, and 1 mM TCEP. Samples were subjected to analytical ultracentrifugation. Samples were centrifuged at 50,000 rpm and monitored at a wavelength of 290 nm. The continuous sedimentation coefficient c(s) was plotted as a function of the sedimentation coefficient (S). (C) ATP consumption by *Pf*AspRS*, Pv*AspRS and *Hs*AspRS in the presence and absence of the *Ec*tRNA. Reagent concentrations: 50 nM *Pf*AspRS and *Pv*AspRS or 100 nM *Hs*AspRS with 10 μM ATP, 200 μM Asp, 1 U/mL pyrophosphatase, 80 μM *Ec*tRNA for *Pf*AspRS and *Pv*AspRS and 160 μM *Ec*tRNA for *Hs*AspRS. Data represent mean ± SEM from four independent experiments. (D-F) Thermal stabilization of native length AspRS enzymes by Asp-DACM. First derivatives of melting curves for native length *Pf*AspRS (D), *Pv*AspRS (E) or *Hs*AspRS (F) (1.5 μM) after incubation at 37°C for 3 h with 2 or 5 μM Asp-DACM. Data are representative of three independent experiments.

Supplementary Figure 6. Structure prediction for native length *Pf*AspRS and *Pv*AspRS and biochemical analysis of truncated *Pv*AspRS.

(A) AlphaFold predicted structures of native length *Pf*AspRS (PlasmoDB ID: PF3D7_0102900) and native length *Pv*AspRS (PlasmoDB ID: PVX_081610). Model confidence is predicted and colored. Blue: Very high (pLDDT > 90), sky blue (90 > pLDDT > 70), yellow (70 > pLDDT > 50), and orange: very low (pLDDT < 50) per-residue model confidence score (pLDDT). (B) ATP consumption by *Pv*AspRS (51-631) and *Pv*AspRS (96-631) in the presence and absence of the *Ec*tRNA. 50 nM *Pv*AspRS was incubated with 10 μM ATP, 200 μM Asp, 1 U/mL pyrophosphatase, ± 80 μM EctRNA in 25 mM Tris-HCl (pH 8), 150 mM NaCl, 5 mM MgCl_2_, 1 mM TCEP, 0.1 mg/mL BSA for 1 h at 37℃. Data represent five independent experiments. (C,D) First derivatives of melting curves for native length *Pf*AspRS (1.5 μM) in apo form or after incubation at 37°C for 3 h with 10 μM ATP, 20 μM Asp, 80 μM *Ec*tRNA, 10-50 μM DACM (C) or 2-10 μM Asp-DACM (D). Data are representative of three independent experiments.

Supplementary Figure 7. Comparison of crystal structure of Asp-DACM-bound *Pv*AspRS with apo *Pv*AspRS.

(A) Overlay of chain A of apo *Pv*AspRS and Asp-DACM-bound *Pv*AspRS showing the flipping loop and the motif II loop. (B) Ribbon representation of the flipping loop of chain B of apo *Pv*AspRS. The SEGG loop interacts with a conserved residue Q371, as well as N356, S350 and A357. (C) B-factor analysis of the chain A and B flipping loops of different *Pv*AspRS structures. The x-axis shows residue number.

Supplementary Figure 8. Crystal structure of Asp-AMP-bound *Pv*AspRS and comparison with the apo and Asp-DACM-bound structures.

(A) 2*F*_o_-*F*_c_ maps contoured at 2 σ (mesh surface) showing electron density supporting the position of Asp-AMP bound to chains A and B. (B) Ligplots of Asp-AMP active site interfaces for the A and B chains. Hydrogen bonds and salt bridges are depicted with dashed (green) lines. Other interactions between protein and ligand are indicated by red arcs. (C) Overlay of *Pv*AspRS (Apo), *Pv*AspRS (Asp-DACM) and *Pv*AspRS (Asp-AMP) showing the flipping loop and the motif II loop. Chain A (left), Chain B (right). The sidechain of residue E398 of apo *Pv*AspRS Chain A is not built due to insufficient density.

---

**Supplementary Table 1.** Thermal stabilization of recombinant *Pf*TyrRS, native length *Pf, Pv, Hs*AspRS and truncated *Pv*AspRS by nucleoside sulfamates. *N.D. = Not determined due to incomplete transition.

|  |  | **T<sub>m</sub><sub>app</sub> (°C)**<br>**Mean ± SEM** | **Δ T<sub>m</sub><sub>app</sub>**<br>**(°C)** | **n** |
| --- | --- | --- | --- | --- |
| ***Pf*TyrRS** | *Pf*TyrRS alone | 49.3 ± 0.0 | - | 3 |
|  | *Pf*TyrRS + ATP + Asp + *Pf*tRNA(Tyr) | 50.4 ± 0.1 | - | 3 |
|  | *Pf*TyrRS + ATP + Asp + *Pf*tRNA(Tyr) + 5 μM DACM | 50.0 ± 0.1 | -0.3 | 3 |
|  | *Pf*TyrRS + ATP + Asp + *Pf*tRNA(Tyr) + 10 μM DACM | 50.8 ± 0.6 | 0.4 | 3 |
|  | *Pf*TyrRS + ATP + Asp + *Pf*tRNA(Tyr) + 5 μM AMS | N.D.* | N.D.* | 3 |
|  | *Pf*TyrRS + ATP + Asp + t *Pf*tRNA(Tyr) + 10 μM AMS | 60.6 ± 0.4 | 10.2 | 3 |
| ***Pf*AspRS**<br>**(49-626)** | *Pf*AspRS alone | 47.3 ± 0.2 | - | 3 |
|  | *Pf*AspRS + ATP + Asp + *Ect*RNA | 43.0 ± 0.5 | - | 6 |
|  | *Pf*AspRS + ATP + Asp + *Ect*RNA + 10 μM DACM | N.D.* | N.D.* | 3 |
|  | *Pf*AspRS + ATP + Asp + *Ect*RNA + 20 μM DACM | N.D.* | N.D.* | 3 |
|  | *Pf*AspRS + ATP + Asp + *Ect*RNA + 50 μM DACM | 61.1 ± 0.1 | 18.1 | 3 |
|  | *Pf*AspRS + 2 μM Asp-DACM | N.D.* | N.D.* | 3 |
|  | *Pf*AspRS + 5 μM Asp-DACM | 65.3 ± 0.1 | 18.0 | 3 |
|  | *Pf*AspRS + 10 μM Asp-DACM | 65.7 ± 0.1 | 18.4 | 3 |
|  | *Pf*AspRS + ATP + Asp + *Ect*RNA + 10 μM AMS | N.D.* | N.D.* | 3 |
|  | *Pf*AspRS + ATP + Asp + *Ect*RNA + 20 μM AMS | 56.4 ± 0.2 | 13.4 | 3 |
|  | *Pf*AspRS + ATP + Asp + *Ect*RNA + 50 μM AMS | 57.0 ± 0.2 | 14.0 | 3 |
| ***Pv*AspRS**<br>**(51-631)** | *Pv*AspRS alone | 46.8 ± 0.03 | - | 3 |
|  | *Pv*AspRS + ATP + Asp + *Ect*RNA | 43.4 ± 0.3 | - | 6 |
|  | *Pv*AspRS + ATP + Asp + *Ect*RNA + 10 μM DACM | 62.4 ± 0.02 | 19.0 | 3 |
|  | *Pv*AspRS + ATP + Asp + *Ect*RNA + 20 μM DACM | 62.6 ± 0.2 | 19.2 | 3 |
|  | *Pv*AspRS + ATP + Asp + *Ect*RNA + 50 μM DACM | 62.6 ± 0.2 | 19.2 | 3 |
|  | *Pv*AspRS + 2 μM Asp-DACM | N.D.* | N.D.* | 3 |
|  | *Pv*AspRS + 5 μM Asp-DACM | 66.3 ± 0.005 | 19.5 | 3 |
|  | *Pv*AspRS + 10 μM Asp-DACM | 66.8 ± 0.05 | 20.0 | 3 |
| | *PvAspRS* + ATP + Asp + *EctRNA* + 10 μM AMS | 59.3 ± 0.3 | 15.9 | 3 |
| | *PvAspRS* + ATP + Asp + *EctRNA* + 20 μM AMS | 59.3 ± 0.2 | 15.9 | 3 |
| | *PvAspRS* + ATP + Asp + *EctRNA* + 50 μM AMS | 59.5 ± 0.2 | 16.1 | 3 |
| ***HsAspRS***<br>(1-501) | *HsAspRS* alone | 52.1 ± 0.02 | - | 3 |
| | *HsAspRS* + ATP + Asp + *EctRNA* | 49.1 ± 0.2 | - | 6 |
| | *HsAspRS* + ATP + Asp + *EctRNA* + 10 μM DACM | N.D.* | N.D.* | 3 |
| | *HsAspRS* + ATP + Asp + *EctRNA* + 20 μM DACM | N.D.* | N.D.* | 3 |
| | *HsAspRS* + ATP + Asp + *EctRNA* + 50 μM DACM | N.D.* | N.D.* | 3 |
| | *HsAspRS* + 2 μM Asp-DACM | N.D.* | N.D.* | 3 |
| | *HsAspRS* + 5 μM Asp-DACM | 75.5 ± 0.1 | 23.4 | 3 |
| | *HsAspRS* + 10 μM Asp-DACM | 76.0 ± 0.2 | 23.9 | 3 |
| | *HsAspRS* + ATP + Asp + *EctRNA* + 10 μM AMS | 48.7 ± 0.3 | -0.4 | 3 |
| | *HsAspRS* + ATP + Asp + *EctRNA* + 20 μM AMS | 48.6 ± 0.1 | -0.5 | 3 |
| | *HsAspRS* + ATP + Asp + *EctRNA* + 50 μM AMS | 48.7 ± 0.2 | -0.4 | 3 |
| ***PvAspRS***<br>(96-631) | *PvAspRS* (96-631) alone | 46.3 ± 0.2 | - | 3 |
| | *PfAspRS* (96-631) + ATP + Asp + *EctRNA* | 42.5 ± 0.2 | - | 3 |
| | *PfAspRS* (96-631)+ ATP + Asp + *EctRNA* + 10 μM DACM | N.D.* | N.D.* | 3 |
| | *PfAspRS* (96-631)+ ATP + Asp + *EctRNA* + 20 μM DACM | N.D.* | N.D.* | 3 |
| | *PfAspRS* (96-631)+ ATP + Asp + *EctRNA* + 50 μM DACM | 60.9 ± 0.1 | 18.4 | 3 |
| | *PvAspRS* + 2 μM Asp-DACM | 64.2 ± 0.1 | 17.9 | 3 |
| | *PvAspRS* + 5 μM Asp-DACM | 66.0 ± 0.2 | 19.7 | 3 |
| | *PvAspRS* + 10 μM Asp-DACM | 66.4 ± 0.0 | 20.1 | 3 |

**Supplementary Table 2.** Protein–ligand docking scores determined using the Surflex fragment matching strategy.

| Compound | Tyrosine fragment docking score |
| --- | --- |
| Tyr-AMP | 15.5 |
| Tyr-AMS | 15.0 |
| Tyr-DACM | 13.6 |

**Supplementary Table 3.**
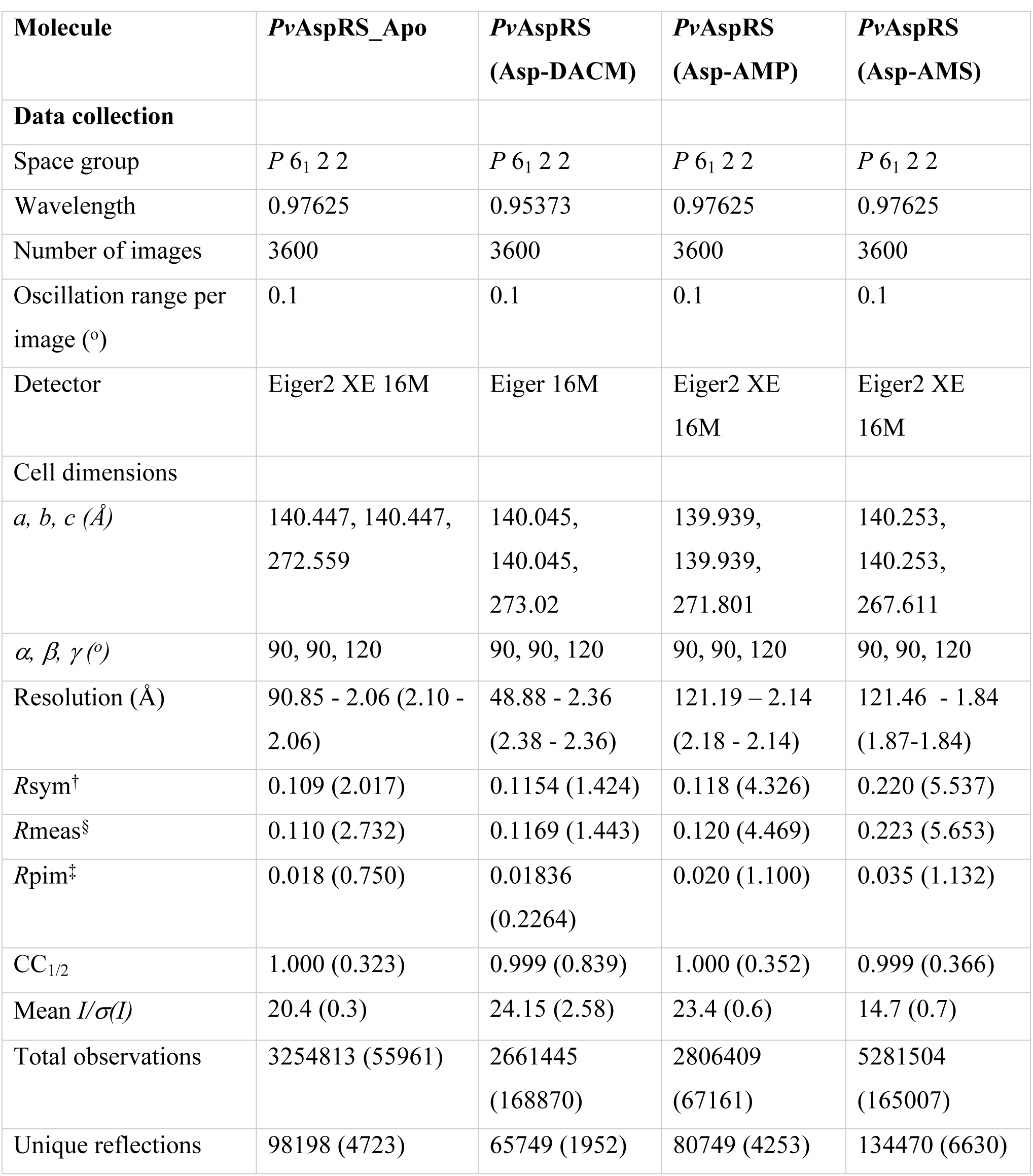
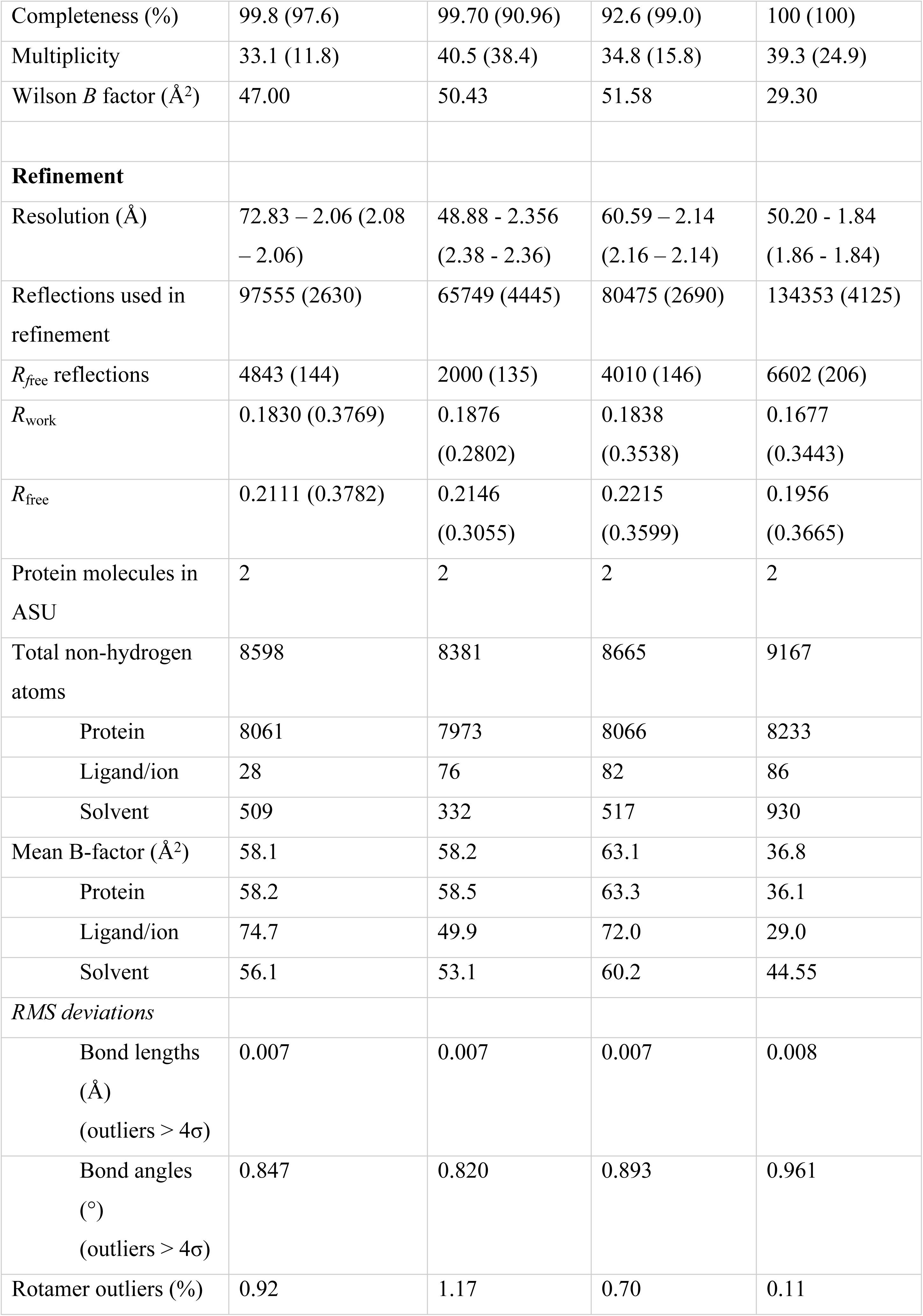
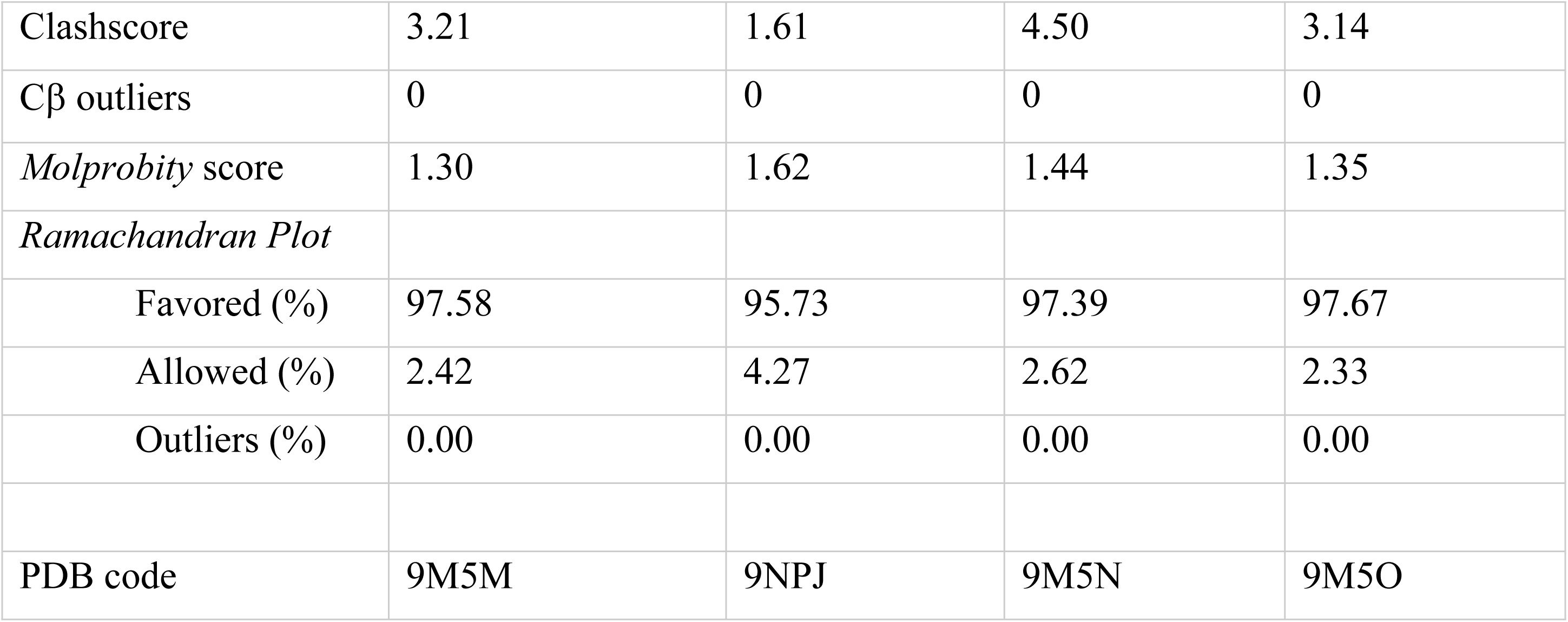
X-ray diffraction data collection and refinement statistics for truncated apo *Pv*AspRS and in complex with ligands Asp-DACM, Asp-AMP and Asp-AMS. Values for the highest resolution shell are given in parentheses. ^†^ *R*sym = ΣhklΣi |Ii (hkl) - 〈I(hkl)〉| / ΣhklΣiIi (hkl) ^§^ *R*meas= Σhkl [N/(N - 1)]^½^ Σi |Ii (hkl) - 〈I (hkl)〉| / Σhkl Σi Ii (hkl) ^‡^ *R*pim= Σhkl [1/(N - 1)]^½^ Σi |Ii (hkl) - 〈I (hkl)〉| / Σhkl Σi Ii (hkl) *CC*_1/2_ = Pearson correlation coefficient between independently merged half data sets.

## References

1. World_Health_Organisation. WHO World Malaria Report 2024. Geneva: World Health Organization; Licence: CC BY-NC-SA 30 IGO. 2024.

2. Knox TB, Juma EO, Ochomo EO, Pates Jamet H, Ndungo L, Chege P, et al. An online tool for mapping insecticide resistance in major Anopheles vectors of human malaria parasites and review of resistance status for the Afrotropical region. Parasites & Vectors. 2014;7(1):76.

3. Flegg JA, Kandanaarachchi S, Guerin PJ, Dondorp AM, Nosten FH, Otienoburu SD, et al. Spatio-temporal spread of artemisinin resistance in Southeast Asia. PLoS computational biology. 2024;20(4):e1012017.

4. Hamilton WL, Amato R, van der Pluijm RW, Jacob CG, Quang HH, Thuy-Nhien NT, et al. Evolution and expansion of multidrug-resistant malaria in southeast Asia: a genomic epidemiology study. The Lancet Infectious diseases. 2019;19(9):943–51.

5. Rosenthal PJ, Asua V, Conrad MD. Emergence, transmission dynamics and mechanisms of artemisinin partial resistance in malaria parasites in Africa. Nature Reviews Microbiology. 2024;22(6):373–84.

6. Dhorda M, Kaneko A, Komatsu R, Kc A, Mshamu S, Gesase S, et al. Artemisinin-resistant malaria in Africa demands urgent action. Science. 2024;385(6706):252–4.

7. Francklyn CS, Mullen P. Progress and challenges in aminoacyl-tRNA synthetase-based therapeutics. J Biol Chem. 2019;294(14):5365–85.

8. Xie SC, Griffin MDW, Winzeler EA, Ribas de Pouplana L, Tilley L. Targeting Aminoacyl tRNA Synthetases for Antimalarial Drug Development. Annu Rev Microbiol. 2023;77:111–29.

9. Gill J, Sharma A. Exploration of aminoacyl-tRNA synthetases from eukaryotic parasites for drug development. J Biol Chem. 2023;299(3):102860.

10. Khoshnood S, Heidary M, Asadi A, Soleimani S, Motahar M, Savari M, et al. A review on mechanism of action, resistance, synergism, and clinical implications of mupirocin against Staphylococcus aureus. Biomed Pharmacother. 2019;109:1809–18.

11. Pines M, Spector I. Halofuginone - the multifaceted molecule. Molecules. 2015;20(1):573–94.

12. Hyer ML, Milhollen MA, Ciavarri J, Fleming P, Traore T, Sappal D, et al. A small-molecule inhibitor of the ubiquitin activating enzyme for cancer treatment. Nat Med. 2018;24(2):186–93.

13. Brownell JE, Sintchak MD, Gavin JM, Liao H, Bruzzese FJ, Bump NJ, et al. Substrate-assisted inhibition of ubiquitin-like protein-activating enzymes: the NEDD8 E1 inhibitor MLN4924 forms a NEDD8-AMP mimetic in situ. Mol Cell. 2010;37(1):102–11.

14. Langston SP, Grossman S, England D, Afroze R, Bence N, Bowman D, et al. Discovery of TAK-981, a first-in-class inhibitor of SUMO-activating enzyme for the treatment of cancer. Journal of medicinal chemistry. 2021;64(5):2501–20.

15. Xie SC, Metcalfe RD, Dunn E, Morton CJ, Huang SC, Puhalovich T, et al. Reaction hijacking of tyrosine tRNA synthetase as a new whole-of-life-cycle antimalarial strategy. Science. 2022;376(6597):1074–9.

16. Xie SC, Tai C-W, Morton CJ, Ma L, Huang S-C, Wittlin S, et al. A potent and selective reaction hijacking inhibitor of *Plasmodium falciparum* tyrosine tRNA synthetase exhibits single dose oral efficacy in vivo. PLoS pathogens. 2024;20(12):e1012429.

17. Xie SC, Wang Y, Morton CJ, Metcalfe RD, Dogovski C, Pasaje CFA, et al. Reaction hijacking inhibition of *Plasmodium falciparum* asparagine tRNA synthetase. Nat Commun. 2024;15(1):937.

18. Scacchi A, Bortolo R, Cassani G, Nielsen E. Herbicidal Activity of Dealanylascamycin, a Nucleoside Antibiotic. Pesticide Biochemistry and Physiology. 1994;50(2):149–58.

19. Takahashi E, Beppu T. A new nucleosidic antibiotic AT-265. J Antibiot (Tokyo). 1982;35(8):939–47.

20. Osada H, Isono K. Mechanism of action and selective toxicity of ascamycin, a nucleoside antibiotic. Antimicrobial agents and chemotherapy. 1985;27(2):230–3.

21. Awakawa T, Barra L, Abe I. Biosynthesis of sulfonamide and sulfamate antibiotics in actinomycete. J Ind Microbiol Biotechnol. 2021;48(3-4).

22. Bloch A, Coutsogeorgopoulos C. Inhibition of protein synthesis by 5’-sulfamoyladenosine. Biochemistry. 1971;10(24):4395–8.

23. Mujumdar P, Poulsen SA. Natural product primary sulfonamides and primary sulfamates. Journal of natural products. 2015;78(6):1470–7.

24. Mujumdar P, Bua S, Supuran CT, Peat TS, Poulsen SA. Synthesis, structure and bioactivity of primary sulfamate-containing natural products. Bioorg Med Chem Lett. 2018;28(17):3009–13.

25. Voorneveld J, Rack JGM, van Gijlswijk L, Meeuwenoord NJ, Liu Q, Overkleeft HS, et al. Molecular tools for the study of ADP-ribosylation: A Unified and versatile method to synthesise native mono-ADP-ribosylated peptides. Chemistry – A European Journal. 2021;27(41):10621–7.

26. Castro-Pichel J, García-López MT, De las Heras FG. A facile synthesis of ascamycin and related analogues. Tetrahedron. 1987;43(2):383–9.

27. Cain R, Salimraj R, Punekar AS, Bellini D, Fishwick CWG, Czaplewski L, et al. Structure-guided enhancement of selectivity of chemical robe inhibitors targeting bacterial seryl-tRNA synthetase. J Med Chem. 2019;62(21):9703–17.

28. Fu Y, Tilley L, Kenny S, Klonis N. Dual labeling with a far red probe permits analysis of growth and oxidative stress in *P. falciparum*-infected erythrocytes. Cytometry A. 2010;77(3):253–63.

29. Solyakov L, Halbert J, Alam MM, Semblat JP, Dorin-Semblat D, Reininger L, et al. Global kinomic and phospho-proteomic analyses of the human malaria parasite *Plasmodium falciparum*. Nat Commun. 2011;2:565.

30. Bridgford JL, Xie SC, Cobbold SA, Pasaje CFA, Herrmann S, Yang T, et al. Artemisinin kills malaria parasites by damaging proteins and inhibiting the proteasome. Nature Communications. 2018;9(1):3801.

31. Liu J, Xu Y, Stoleru D, Salic A. Imaging protein synthesis in cells and tissues with an alkyne analog of puromycin. Proc Natl Acad Sci U S A. 2012;109(2):413–8.

32. Du Y, Giannangelo C, He W, Shami GJ, Zhou W, Yang T, et al. Dimeric artesunate glycerophosphocholine conjugate nano-assemblies as slow-release antimalarials to overcome Kelch 13 mutant artemisinin resistance. Antimicrob Agents Chemother. 2022;66(5):e0206521.

33. Jain AN. Surflex: Fully automatic flexible molecular docking using a molecular similarity-based search engine. Journal of medicinal chemistry. 2003;46(4):499–511.

34. Eriani G, Delarue M, Poch O, Gangloff J, Moras D. Partition of tRNA synthetases into two classes based on mutually exclusive sets of sequence motifs. Nature. 1990;347(6289):203-6.

35. Bour T, Akaddar A, Lorber B, Blais S, Balg C, Candolfi E, et al. Plasmodial aspartyl-tRNA synthetases and peculiarities in *Plasmodium falciparum*. J Biol Chem. 2009;284(28):18893–903.

36. Frugier M, Moulinier L, Giegé R. A domain in the N-terminal extension of class IIb eukaryotic aminoacyl-tRNA synthetases is important for tRNA binding. Embo J. 2000;19(10):2371–80.

37. Abbott JA, Livingston NM, Egri SB, Guth E, Francklyn CS. Characterization of aminoacyl-tRNA synthetase stability and substrate interaction by differential scanning fluorimetry. Methods (San Diego, Calif). 2017;113:64–71.

38. Moulinier L, Eiler S, Eriani G, Gangloff J, Thierry JC, Gabriel K, et al. The structure of an AspRS-tRNA(Asp) complex reveals a tRNA-dependent control mechanism. Embo J. 2001;20(18):5290–301.

39. Eiler S, Dock-Bregeon A, Moulinier L, Thierry JC, Moras D. Synthesis of aspartyl-tRNA(Asp) in *Escherichia coli* - a snapshot of the second step. Embo J. 1999;18(22):6532–41.

40. Kim KR, Park SH, Kim HS, Rhee KH, Kim BG, Kim DG, et al. Crystal structure of human cytosolic aspartyl-tRNA synthetase, a component of multi-tRNA synthetase complex. Proteins. 2013;81(10):1840–6.

41. Cavarelli J, Rees B, Ruff M, Thierry JC, Moras D. Yeast tRNA(Asp) recognition by its cognate class II aminoacyl-tRNA synthetase. Nature. 1993;362(6416):181-4.

42. Eriani G, Cavarelli J, Martin F, Ador L, Rees B, Thierry JC, et al. The class II aminoacyl-tRNA synthetases and their active site: evolutionary conservation of an ATP binding site. J Mol Evol. 1995;40(5):499–508.

43. Rees B, Webster G, Delarue M, Boeglin M, Moras D. Aspartyl tRNA-synthetase from Escherichia coli: flexibility and adaptability to the substrates. J Mol Biol. 2000;299(5):1157–64.

44. Thomas SO, Singleton VL, Lowery JA, Sharpe RW, Pruess LM, Porter JN, et al. Nucleocidin, a new antibiotic with activity against Trypanosomes. Antibiot Annu. 1956:716–21.

45. Waller CW, Patrick JB, Fulmor W, Meyer WE. The structure of nucleocidin. I. Journal of the American Chemical Society. 1957;79(4):1011–2.

46. Isono K, Uramoto M, Kusakabe H, Miyata N, Koyama T, Ubukata M, et al. Ascamycin and dealanylascamycin, nucleoside antibiotics from Streptomyces sp. J Antibiot (Tokyo). 1984;37(6):670–2.

47. Florini JR, Bird HH, Bell PH. Inhibition of protein synthesis in vitro and in vivo by nucleocidin, an antitrypanosomal antibiotic. J Biol Chem. 1966;241(5):1091–8.

48. Castilho BA, Shanmugam R, Silva RC, Ramesh R, Himme BM, Sattlegger E. Keeping the eIF2 alpha kinase Gcn2 in check. Biochim Biophys Acta. 2014;1843(9):1948–68.

49. Fennell C, Babbitt S, Russo I, Wilkes J, Ranford-Cartwright L, Goldberg DE, et al. PfeIK1, a eukaryotic initiation factor 2alpha kinase of the human malaria parasite *Plasmodium falciparum*, regulates stress-response to amino-acid starvation. Malar J. 2009;8:99.

50. Reed VS, Yang DC. Characterization of a novel N-terminal peptide in human aspartyl-tRNA synthetase. Roles in the transfer of aminoacyl-tRNA from aminoacyl-tRNA synthetase to the elongation factor 1 alpha. J Biol Chem. 1994;269(52):32937–41.

51. Cheong H-K, Park J-Y, Kim E-H, Lee C, Kim S, Kim Y, et al. Structure of the N-terminal extension of human aspartyl-tRNA synthetase: implications for its biological function. The international journal of biochemistry & cell biology. 2003;35(11):1548–57.

52. Schmitt E, Moulinier L, Fujiwara S, Imanaka T, Thierry JC, Moras D. Crystal structure of aspartyl-tRNA synthetase from *Pyrococcus kodakaraensis* KOD: archaeon specificity and catalytic mechanism of adenylate formation. Embo J. 1998;17(17):5227–37.

53. Sauter C, Lorber B, Cavarelli J, Moras D, Giege R. The free yeast aspartyl-tRNA synthetase differs from the tRNA(Asp)-complexed enzyme by structural changes in the catalytic site, hinge region, and anticodon-binding domain. J Mol Biol. 2000;299(5):1313–24.

54. Lawrence G, Cheng QQ, Reed C, Taylor D, Stowers A, Cloonan N, et al. Effect of vaccination with 3 recombinant asexual-stage malaria antigens on initial growth rates of *Plasmodium falciparum* in non-immune volunteers. Vaccine. 2000;18(18):1925–31.

55. Donato MT, Tolosa L, Gómez-Lechón MJ. Culture and functional characterization of human hepatoma HepG2 cells. Methods Mol Biol. 2015;1250:77–93.

56. Straimer J, Gnadig NF, Witkowski B, Amaratunga C, Duru V, Ramadani AP, et al. Drug resistance. K13-propeller mutations confer artemisinin resistance in *Plasmodium falciparum* clinical isolates. Science. 2015;347(6220):428–31.

57. Schuck P. Size-distribution analysis of macromolecules by sedimentation velocity ultracentrifugation and lamm equation modeling. Biophys J. 2000;78(3):1606–19.

58. Rio DC, Ares M, Jr., Hannon GJ, Nilsen TW. Purification of RNA using TRIzol (TRI reagent). Cold Spring Harb Protoc. 2010;2010(6):pdb prot5439.

59. Gorrec F. The MORPHEUS II protein crystallization screen. Acta Crystallogr F Struct Biol Commun. 2015;71(Pt 7):831–7.

60. Aragão D, Aishima J, Cherukuvada H, Clarken R, Clift M, Cowieson NP, et al. MX2: a high-flux undulator microfocus beamline serving both the chemical and macromolecular crystallography communities at the Australian Synchrotron. J Synchrotron Radiat. 2018;25(Pt 3):885–91.

61. Kabsch W. XDS. Acta Crystallogr D Biol Crystallogr. 2010;66(Pt 2):125–32.

62. Evans PR, Murshudov GN. How good are my data and what is the resolution? Acta Crystallogr D Biol Crystallogr. 2013;69(Pt 7):1204–14.

63. Winter G, Waterman DG, Parkhurst JM, Brewster AS, Gildea RJ, Gerstel M, et al. DIALS: implementation and evaluation of a new integration package. Acta Crystallogr D Struct Biol. 2018;74(Pt 2):85–97.

64. Vonrhein C, Flensburg C, Keller P, Sharff A, Smart O, Paciorek W, et al. Data processing and analysis with the autoPROC toolbox. Acta Crystallogr D Biol Crystallogr. 2011;67(Pt 4):293–302.

65. McCoy AJ, Grosse-Kunstleve RW, Adams PD, Winn MD, Storoni LC, Read RJ. Phaser crystallographic software. J Appl Crystallogr. 2007;40(Pt 4):658–74.

66. Panjikar S, Venkataraman P, Lamzin V, Weiss M, Tucker P. Auto-Rickshaw: an automated crystal structure determination platform as an efficient tool for the validation of an X-ray diffraction experiment. Acta Crystallogr D Biol Crystallogr. 2005;61:449–57.

67. Afonine PV, Grosse-Kunstleve RW, Echols N, Headd JJ, Moriarty NW, Mustyakimov M, et al. Towards automated crystallographic structure refinement with phenix.refine. Acta Crystallogr D Biol Crystallogr. 2012;68(Pt 4):352–67.

68. Liebschner D, Afonine PV, Baker ML, Bunkóczi G, Chen VB, Croll TI, et al. Macromolecular structure determination using X-rays, neutrons and electrons: recent developments in Phenix. Acta Crystallogr D Struct Biol. 2019;75(Pt 10):861–77.

69. Emsley P, Lohkamp B, Scott WG, Cowtan K. Features and development of Coot. Acta Crystallogr D Biol Crystallogr. 2010;66(Pt 4):486–501.

70. Moriarty NW, Grosse-Kunstleve RW, Adams PD. electronic Ligand Builder and Optimization Workbench (eLBOW): a tool for ligand coordinate and restraint generation. Acta Crystallogr D Biol Crystallogr. 2009;65(Pt 10):1074–80.

71. Chen VB, Arendall WB, 3rd, Headd JJ, Keedy DA, Immormino RM, Kapral GJ, et al. MolProbity: all-atom structure validation for macromolecular crystallography. Acta Crystallogr D Biol Crystallogr. 2010;66(Pt 1):12–21.

72. Williams CJ, Headd JJ, Moriarty NW, Prisant MG, Videau LL, Deis LN, et al. MolProbity: More and better reference data for improved all-atom structure validation. Protein Sci. 2018;27(1):293–315.

73. Pettersen EF, Goddard TD, Huang CC, Couch GS, Greenblatt DM, Meng EC, et al. UCSF Chimera - a visualization system for exploratory research and analysis. J Comput Chem. 2004;25(13):1605–12.

74. Zheng H, Cooper DR, Porebski PJ, Shabalin IG, Handing KB, Minor W. CheckMyMetal: a macromolecular metal-binding validation tool. Acta Crystallogr D Struct Biol. 2017;73(Pt 3):223–33.

75. Laskowski RA, Swindells MB. LigPlot+: Multiple ligand–protein interaction diagrams for drug discovery. Journal of Chemical Information and Modeling. 2011;51(10):2778–86.

76. Schrödinger L. The PyMOL Molecular Graphics System, Version 2.5.4.

77. Meng EC, Goddard TD, Pettersen EF, Couch GS, Pearson ZJ, Morris JH, et al. UCSF ChimeraX: Tools for structure building and analysis. Protein Science. 2023;32(11):e4792.

